# Local frustration determines loop opening during the catalytic cycle of an oxidoreductase

**DOI:** 10.1101/686949

**Authors:** Lukas L. Stelzl, Despoina A.I. Mavridou, Emmanuel Saridakis, Diego Gonzalez, Andrew J. Baldwin, Stuart J. Ferguson, Mark S.P. Sansom, Christina Redfield

**Affiliations:** Department of Biochemistry, University of Oxford, South Parks Road, Oxford OX1 3QU, United Kingdom; Department of Molecular Biosciences, University of Texas at Austin, Austin, Texas, 78712, U.S.A; Institute of Nanoscience and Nanotechnology, N.C.S.R. Demokritos, Aghia Paraskevi, Athens 15310, Greece; Laboratoire de Microbiologie, Institut de Biologie, Université de Neuchâtel, Neuchâtel, 2000, Switzerland; Department of Chemistry, Physical and Theoretical Chemistry Laboratory, University of Oxford, South Parks Road, Oxford OX1 3QZ, United Kingdom; Department of Theoretical Biophysics, Max Planck Institute of Biophysics, Max-von-Laue Str. 3, 60438 Frankfurt, Germany

**Keywords:** local frustration, NMR, molecular dynamics, protein dynamics, oxidoreductase

## Abstract

Local structural frustration, the existence of mutually exclusive competing interactions, may explain why some proteins are dynamic while others are rigid. Frustration is thought to underpin biomolecular recognition and the flexibility of protein binding sites. Here we show how a small chemical modification, the oxidation of two cysteine thiols to a disulfide bond, during the catalytic cycle of the N-terminal domain of the key bacterial oxidoreductase DsbD (nDsbD), introduces frustration ultimately influencing protein function. In oxidized nDsbD, local frustration disrupts the packing of the protective cap-loop region against the active site allowing loop opening. By contrast, in reduced nDsbD the cap loop is rigid, always protecting the active-site thiols from the oxidizing environment of the periplasm. Our results point towards an intricate coupling between the dynamics of the active-site cysteines and of the cap loop which modulates the association reactions of nDsbD with its partners resulting in optimized protein function.

## Introduction

Molecular recognition, the specific non-covalent interaction between two or more molecules, underpins all biological processes. During protein-protein association, molecular recognition often depends on conformational changes in one or both of the protein partners. Important insight into the mechanisms of molecular recognition has come from NMR experiments probing dynamics on a range of timescales (Baldwin and Kay 2009, Sekhar et al. 2018). NMR, and especially the relaxation dispersion method, is unique in providing not only kinetic, but also structural information about the conformational changes leading up to protein association (Lange et al. 2008, Sugase, Dyson, and Wright 2007). In some cases, molecular dynamics (MD) simulations have been successfully paired with NMR experiments (Olsson and Noe 2017, Pan et al. 2019, Robustelli, Stafford, and Palmer 2012, Zhang et al. 2016, Wang, Papaleo, and Lindorff-Larsen 2016, Stiller et al. 2019) to provide a more complete picture of the conformational dynamics landscape (Paul et al. 2017, Peters 2012, Robustelli et al. 2013). However, we still do not have a general understanding of the molecular determinants that enable some proteins to access their bound-like conformation before encountering their binding partners in a ‘conformational-selection’ mechanism, while others undergo conformational change only after recognizing their ligands in an ‘induced-fit’ mechanism (Boehr, Nussinov, and Wright 2009).

Local structural frustration (Baldwin, Knowles, et al. 2011, Ferreiro, Komives, and Wolynes 2018, Freiberger et al. 2019), the existence of multiple favorable interactions which cannot be satisfied at the same time, has emerged as an important concept in rationalizing why some regions in proteins undergo conformational changes in the absence of a binding partner (Ferreiro et al. 2007, 2011, Ferreiro, Komives, and Wolynes 2014). In general, globular proteins are overall minimally frustrated and, thus, adopt well-defined native structures. Nonetheless, the existence of a set of competing local interactions can establish a dynamic equilibrium between distinct conformational states. Locally frustrated interactions (Jenik et al. 2012) are indeed more common for residues located in binding interfaces of protein complexes (Ferreiro et al. 2007) and can play an important role in determining which parts of these proteins are flexible. However, very few studies (Danielsson et al. 2013, Das and Plotkin 2013) have analyzed in atomic detail how frustration shapes conformational dynamics and, therefore, can enable protein association. This lack of understanding hampers the exploitation of local frustration, and associated dynamic equilibria, as a design principle for generating optimal protein constructs that could be used in the production of synthetic molecular machines.

To investigate the role of frustration in conformational dynamics, we have used as a model system the N-terminal domain of the transmembrane protein DsbD (nDsbD), a key oxidoreductase in the periplasm of Gram-negative bacteria. DsbD is a three-domain membrane protein (Figure 1A) which acquires two electrons from cytoplasmic thioredoxin and, through a series of thiol-disulfide exchange reactions involving pairs of conserved cysteine residues, transfers these electrons to multiple periplasmic pathways (Figure 1B/C) (Cho and Collet 2013, Depuydt et al. 2009). nDsbD, the last component of the DsbD thiol-disulfide cascade, is essentially the redox interaction hub in the bacterial periplasm; it first interacts with the C-terminal domain of DsbD (cDsbD) to acquire the electrons provided by cytoplasmic thioredoxin and then passes these on to several globular periplasmic proteins, including DsbC, CcmG and DsbG (Figure 1A/B) (Stirnimann et al. 2006). All these thiol-disulfide exchange steps involve close interactions between nDsbD and its binding partners, which are essential so that their functional cysteine pairs can come into proximity leading to electron transfer (Haebel et al. 2002, Mavridou et al. 2011). As nDsbD is essentially a reductase in the oxidative environment of the bacterial periplasm, its interactions with its multiple partners must happen in an optimal manner, making this protein an ideal model system to study.

**Figure 1.**
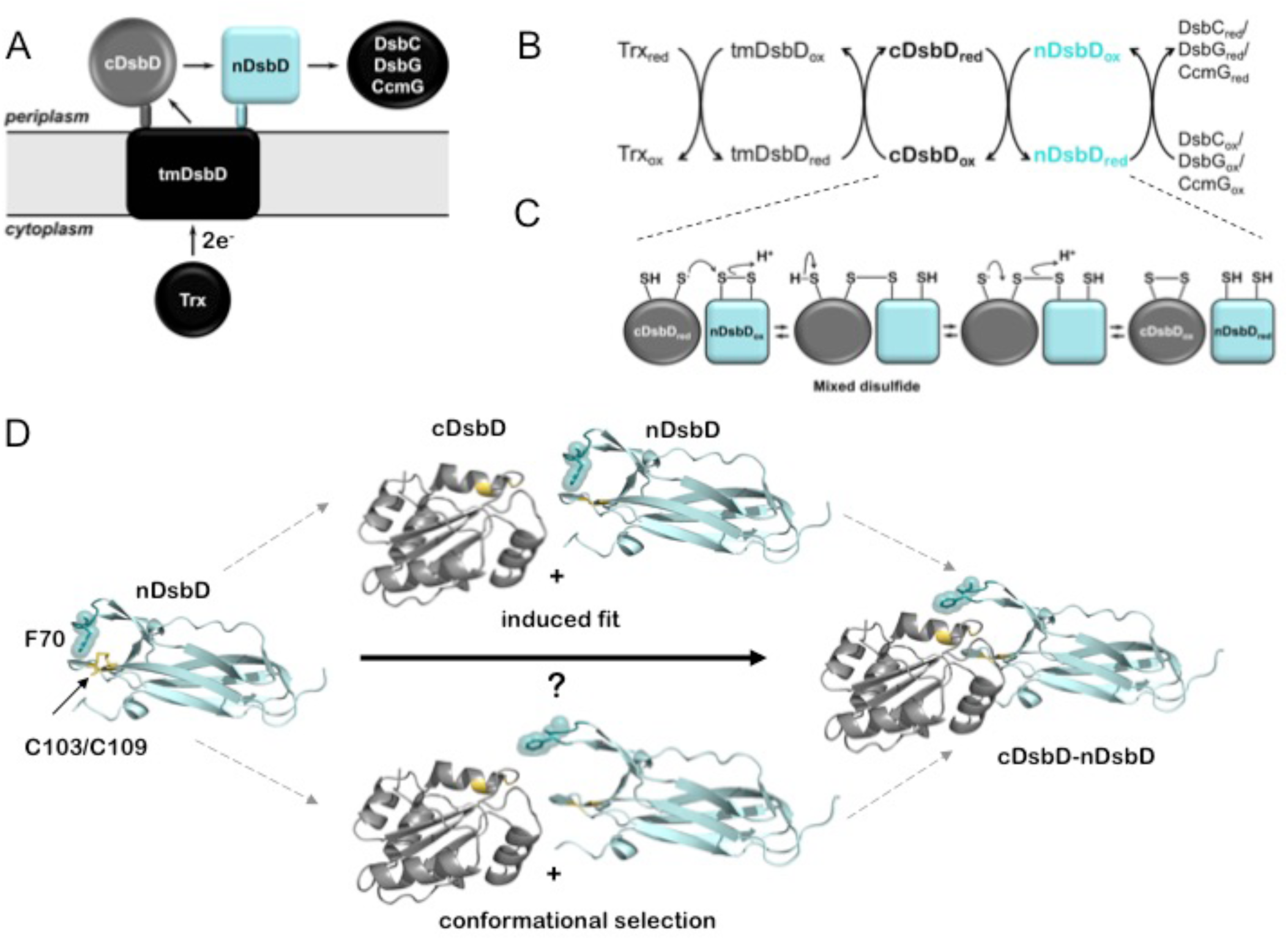
(A) DsbD comprises a central domain with eight transmembrane helices (tmDsbD) and two periplasmic globular domains (nDsbD and cDsbD). The electron transfer pathway from thioredoxin (Trx) to periplasmic partner proteins is shown. cDsbD, DsbC, CcmG and DsbG have a thioredoxin fold (circle), typical of thiol-disulfide oxidoreductases, while nDsbD adopts an immunoglobulin fold (square). nDsbD and cDsbD are shown in cyan and grey, respectively. (B) Electron transfer reactions involving DsbD are shown. Two electrons and two H^+^ are transferred in each step. Trx_red_ reduces the disulfide bond in tmDsbD_ox_, tmDsbD_red_ then reduces cDsbD_ox_, and cDsbD_red_ subsequently reduces nDsbD_ox_. Oxidized DsbC, CcmG and DsbG subsequently accept electrons from nDsbD_red_. (C) Schematic representation of the thiol-disulfide exchange reaction (bimolecular nucleophilic substitution) between cDsbD and nDsbD. (D) The central panels illustrate two possible mechanisms mediating cap loop opening in the formation of the nDsbD-cDsbD complex. In the upper scheme, a steric clash upon binding induces cap loop opening. In the lower scheme, the sampling of an open conformation by nDsbD allows formation of the complex without steric clashes. nDsbD_ox_ is illustrated on the left (PDB: 1L6P); the cap loop (residues 68-72) which covers the active site is in teal. F70 (stick and surface representation) shields the active-site disulfide bond (yellow) in this closed conformation. The nDsbD cap loop adopts an open conformation in the complex between nDsbD and cDsbD (PDB: 1VRS), shown on the right, allowing interaction of the cysteine residues of the two domains. All structures in this and other figures were rendered in Pymol (Version 2.3.5, Schrödinger, LLC) unless otherwise indicated.

X-ray structures show that the cap-loop region of nDsbD, containing residues D68-E69-F70-Y71-G72, plays a key role in the association reactions of this domain with its binding partners (Haebel et al. 2002, Rozhkova et al. 2004, Stirnimann et al. 2005). The catalytic subdomain of nDsbD is inserted at one end of its immunoglobulin (Ig) fold with the two active-site cysteines, C103 and C109, located on opposite strands of a β-hairpin (Figure 1D). For both oxidized and reduced nDsbD (with and without a C103-C109 disulfide bond, respectively) the active-site cysteines are shielded by the cap-loop region which adopts a closed conformation (Figure 1D) (Haebel et al. 2002, Mavridou et al. 2011, Goulding et al. 2002b). Strikingly, all structures of nDsbD in complex with its binding partners, cDsbD, DsbC, and CcmG (Haebel et al. 2002, Rozhkova et al. 2004, Stirnimann et al. 2005), show the cap loop in an open conformation that allows interaction between the cysteine pairs of the two proteins (Figure 1D) (Haebel et al. 2002, Stirnimann et al. 2005, Rozhkova and Glockshuber 2007).

While these X-ray structures provide static snapshots of the closed and open states of the cap loop, they do not offer any insight into the mechanism of loop opening that is essential for the function of nDsbD. To understand the drivers of this conformational change, two hypotheses regarding the solution state of nDsbD need to be examined (Figure 1D). Firstly, it is possible that the cap loop only opens when nDsbD encounters its binding partners, in an ‘induced-fit’ mechanism consistent with the hypothesized protective role of this region of the protein (Goulding et al. 2002b). Secondly, it is also plausible that in solution the cap loop is flexible so that it samples open states, and thus allows binding of partner proteins via ‘conformational selection’ (Boehr, Nussinov, and Wright 2009). Of course, if the cap loop did open in the absence of a binding partner, that could compromise the transfer of electrons in an environment as oxidizing as the periplasm, and particularly through the action of the strong oxidase DsbA (Denoncin and Collet 2013). Therefore, one could hypothesize that a protective role for the cap loop would be more important in reduced nDsbD (nDsbD_red_), raising the question of whether nDsbD interacts with its partners via different loop-opening mechanisms depending on its oxidation state. In this scenario, the pair of cysteines could modulate the conformational ensemble of nDsbD as has been hinted at by structural bioinformatics (Schmidt, Ho, and Hogg 2006, Zhou et al. 2014).

Here we use both state-of-the-art NMR experiments and MD simulations to describe different behaviors for the cap loop depending on the oxidation state of nDsbD. The atomic-scale insight afforded by these two methods reveals how the disulfide bond introduces local frustration specifically in oxidized nDsbD (nDsbD_ox_), allowing the cap loop and active site in nDsbD_ox_ to sample open bound-like conformations in the absence of a binding partner. Our observations have implications not only for the function of DsbD, but more broadly for the role of local frustration in ensuring optimized protein-protein interactions.

## Results and Discussion

### The solution structure of nDsbD as probed by NMR

NMR experiments demonstrate that the cap loop has an important protective role in shielding the active-site cysteines of nDsbD. For nDsbD_ox_, we showed previously that reduction of the disulfide bond by dithiothreitol (DTT) is very slow, with only 50% of nDsbD_ox_ reduced after ∼2 hours. This indicates that the cap loop shields C103 and C109 from the reducing agent (Mavridou et al. 2011). In addition, NMR spectra for nDsbD_red_ collected between pH 6 and 11 show no change in cysteine Cβ chemical shifts. This suggests that the p*K*_a_ of C109, the cysteine residue thought to initiate reductant transfer (Rozhkova et al. 2004), is above 11, thus making it unreactive in isolated nDsbD_red_. This suggests that shielding of the active site by the cap loop in nDsbD_red_ protects C103 and C109 from non-cognate reactions in the oxidizing environment of the periplasm.

The cap loop shields the active site by adopting a closed conformation as in the X-ray structures of oxidized (PDB: 1L6P and 1JPE) (Haebel et al. 2002, Goulding et al. 2002b) and reduced (PDB: 3PFU) (Mavridou et al. 2011) nDsbD. These structures are very similar to each other with pairwise Cα RMSDs for residues 12 to 122 of less than 0.15 Å. To probe the solution structure of oxidized and reduced nDsbD, we measured ^1^H-^15^N residual dipolar couplings (RDCs), which are sensitive reporters of protein structure (Figure 2 – figure supplement 1 and source data 1). Analysis of these data indicates that the orientation of the active-site/cap-loop residues relative to the core β-sandwich of both oxidized and reduced nDsbD in solution is well described by the X-ray structures in which the cap loop adopts a closed conformation. Nevertheless, the cap loop must open to expose the active site in order for nDsbD to carry out its biological function.

### NMR spin-relaxation experiments and model-free analysis

To investigate whether the cap loop is flexible in solution and whether the dynamic behavior of the loop differs in the two oxidation states, we studied the fast time-scale (ps-ns) dynamics of nDsbD using NMR relaxation experiments. ^15^N relaxation experiments, analyzed using the “model-free” approach (Lipari and Szabo 1982a, b), showed that most of the protein backbone is relatively rigid. The majority of order parameters (S^2^) for the backbone amide bonds are above 0.8 (Figure 2). The N- and C-terminal regions of nDsbD gave very low S^2^ values and are clearly disordered in solution. Other residues with order parameters below 0.75 are located in loops or at the start or end of elements of secondary structure, but are not clustered in a specific region. For example, lower order parameters are observed for residues 57 and 58 in both redox states. These residues are located in a long ‘loop’ parallel to the core β-sandwich but not identified as a β-strand due to the absence of hydrogen bonds.

**Figure 2.**
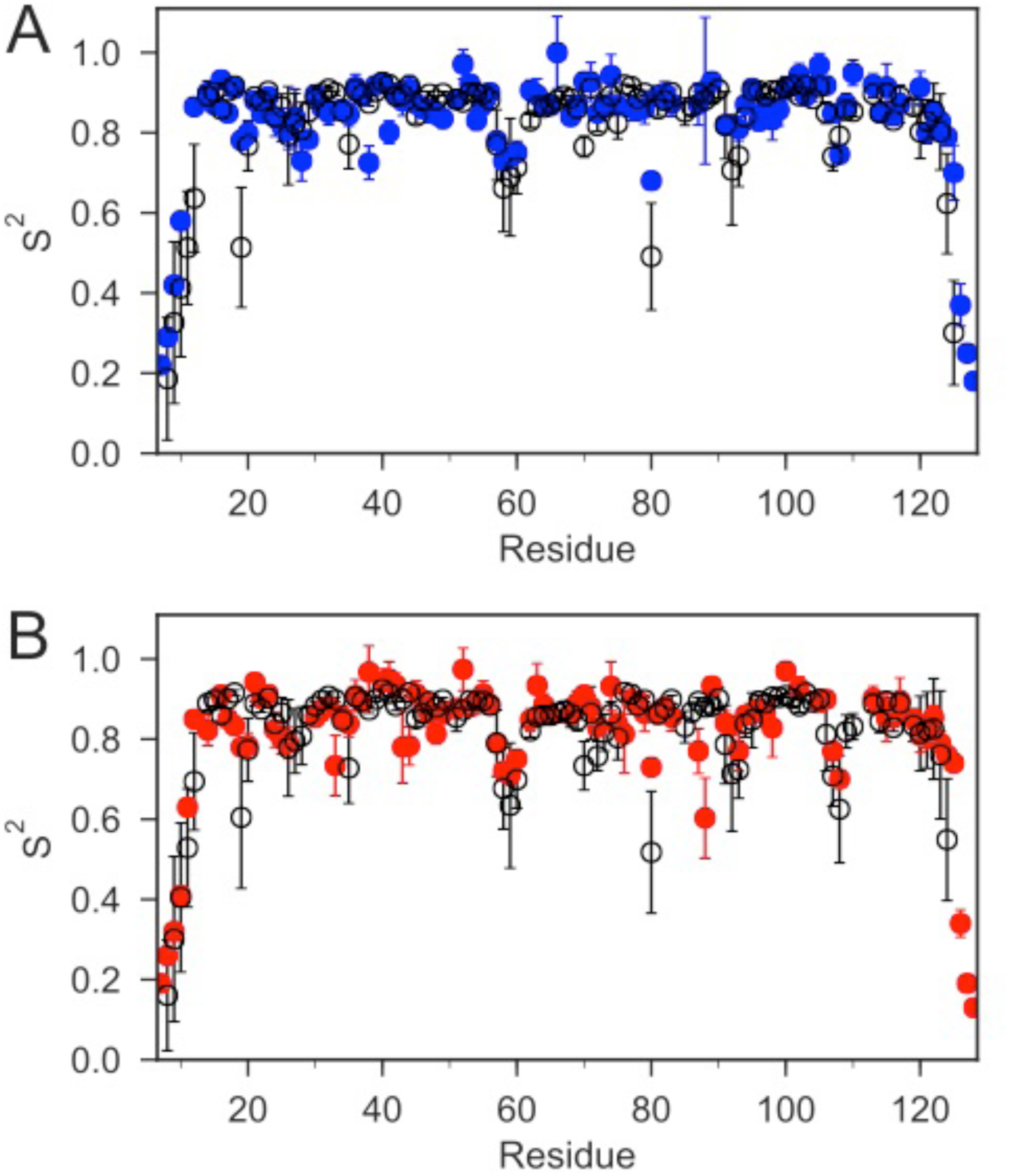
The order parameters (S^2^) for backbone amides derived from ^15^N relaxation experiments and MD simulations are plotted as a function of sequence. The experimental order parameters for reduced and oxidized nDsbD are shown in blue (A) and red (B), respectively. The average order parameters calculated from the combined 1 μs MD simulation ensembles starting from closed structures for reduced (A) and oxidized (B) nDsbD are shown in black. Errors in S^2^ were determined by Monte Carlo simulations (experiment) and a bootstrap analysis (MD simulations). The simulations of both oxidized and reduced nDsbD reproduced even subtle features of sequence-dependent variation of S^2^ values. For example, S^2^ values for residues 57-60 were lowered in both experiment and simulation. These residues are not part of the regular secondary structure of the Ig fold in both the X-ray structures of nDsbD and in the MD simulations.

In both oxidation states, the S^2^ values for the cap-loop region are similar to the rest of the folded protein. Thus, on average, the cap loop of nDsbD does not undergo large amplitude motions on a fast timescale, which means that the active-site cysteines remain shielded. However, close inspection of the {^1^H}-^15^N NOE values for the cap-loop residues in oxidized and reduced nDsbD does show a clear difference between the two oxidation states (Figure 2 – figure supplement 2 and source data 2). In nDsbD_ox_, the residues between V64 and G72 have NOE values at or below 0.7 while in nDsbD_red_ there is only one residue with an NOE value at or below 0.7. The consequence of this difference is that the majority of amino acids in this region of nDsbD_ox_ require a more complex model (with the τ_e_ parameter ranging from ∼50 to 350 ps) to obtain a satisfactory fit in the analysis, in contrast to the majority of residues in nDsbD_red_ can be fitted with the simpler S^2^ only model (with a τ_e_ value of faster than ∼10 ps). Therefore, although the amplitude of motions of cap-loop residues is limited and does not appear to differ very much between the two oxidation states, the timescale of the fast dynamics, as described by the approach of Lipari and Szabo, does differ.

### Molecular dynamics simulations

To evaluate the differences between the conformational dynamics of reduced and oxidized nDsbD with atomic resolution, molecular dynamics (MD) simulations were employed. We tracked the orientation of the cap loop relative to the active site in the generated MD trajectories, by calculating the distance between the center of the aromatic ring of F70 and the sulfur atom of C109. Multiple simulations, for a total of 1 μs, were started from the X-ray structure of reduced nDsbD (3PFU). The distance between F70 and C109 remained close to the value in the X-ray structure of nDsbD_red_, as shown for sample trajectories in Figure 3A and B (Figure 3-figure supplement 2). In addition, C103 and C109 remained largely shielded, as assessed by the solvent accessible surface area (SASA) (Figure 3 – figure supplement 1). Therefore, opening of the cap loop in nDsbD_red_ was not observed in 1 μs of simulation time (Figure 3D and Figure 3 – figure supplement 2). This confirms that the cap loop protects the active-site cysteines of nDsbD_red_ from the oxidizing environment of the periplasm.

**Figure 3.**
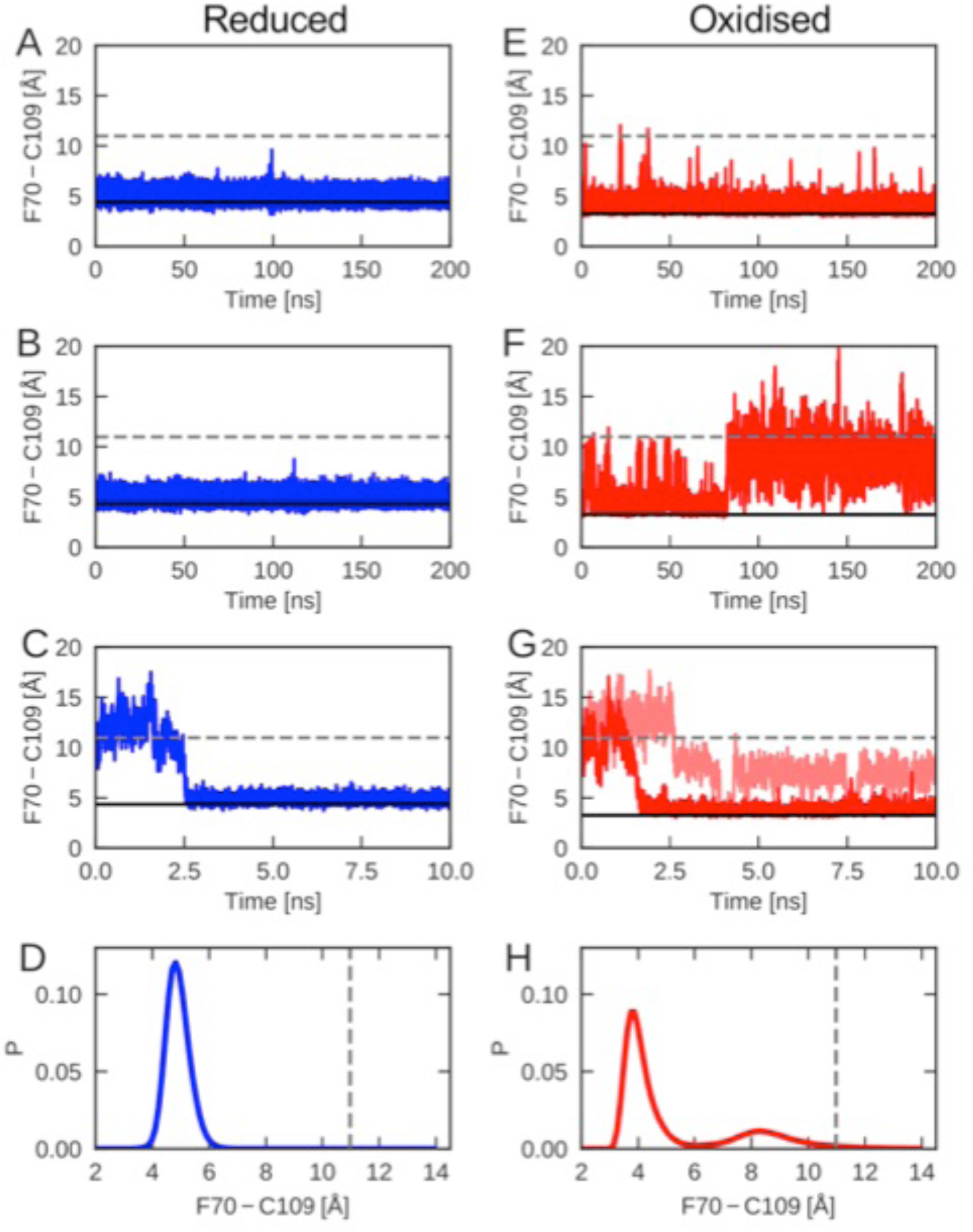
The cap-loop conformation was probed in MD simulations with reduced (A-C) and oxidized (E-G) nDsbD. The conformational state of the cap loop was followed by tracking the distance between the centre of the aromatic ring of F70 and the sulfur atom of C109. The solid and dashed lines at ∼3-4 and ∼11 Å show the F70 ring to C109 Sγ distance in the closed (3PFU/1L6P) and open (1VRS) X-ray structures, respectively. The cap loop remains stably closed in simulations of nDsbD_red_, as shown in (A) and (B). The cap loop in nDsbD_red_ closes within 3 ns for a simulation started from an open conformation based on the 1VRS structure (C). The 1 μs simulation ensemble for nDsbD_red_ shows a closed conformation of F70 relative to C109 (D). The cap loop conformation is more flexible in simulations of nDsbD_ox_; in some simulations it remains closed (E) while in others the loop opens (F). The loop closes in some simulations of nDsbD_ox_ started from an open conformation (G). Overall, the 1 μs simulation ensemble for nDsbD_ox_ shows a preference for the closed conformation of F70 relative to C109, with open conformations sampled relatively rarely (H).

By contrast, nDsbD_ox_ showed more complex dynamics in the 1 μs of simulations launched from the closed 1L6P X-ray structure (Figures 3 and Figure 3 – figure supplement 2). In one 200-ns simulation the cap loop remained closed (Figure 3E), only showing occasional fluctuations. In a different simulation the fluctuations in the cap loop were much more pronounced (Figure 3F); after ∼80 ns the cap loop opened, as judged by the F70-C109 distance, and stayed open for the remainder of the 200 ns MD trajectory. A further 600 ns of simulations showed cap loop opening events with durations ranging from < 1 ns to 22 ns (Figure 3 – figure supplement 2, Figure 5 – figure supplement 4). Calculations of SASA confirmed that opening of the loop increases the solvent exposure of C103 and C109 (Figure 3 – figure supplement 1). The spontaneous loop opening of nDsbD_ox_ and the relative stability of partially open conformations observed in our MD simulations (Figure 3H), mark a clear difference in the conformational ensembles sampled by oxidized nDsbD, as compared to those sampled by the reduced protein (Figure 3D,H).

We have also employed MD to probe the behavior of the cap loop in simulations starting from an open conformation of nDsbD observed in the 1VRS X-ray structure of the nDsbD/cDsbD complex (Rozhkova et al. 2004). Five of the eight trajectories that were started from an open loop conformation for nDsbD_red_, closed within 10 ns (Figure 3C). Similarly, simulations of nDsbD_ox_, that were started from a fully open 1VRS oxidized structure, closed within 10 ns in three out of ten trajectories (Figure 3G). Together, these simulations demonstrate that in both reduced and oxidized nDsbD the cap loop can close rapidly in the absence of a bound interaction partner.

To validate the MD simulations, we used the generated trajectories to predict ^1^H-^15^N RDC and S^2^ values and to compare these to our experimental values. RDCs provide information about the average orientation of peptide bonds and are therefore well-suited for comparison with MD simulations. For both nDsbD_red_ and nDsbD_ox_, the agreement between calculated and experimental RDCs improved for the 1 μs MD ensemble compared to the static X-ray structures (Figure 2 – figure supplement 1 and source data 1). In addition, the extent of the fast-time scale (ps-ns) dynamics observed experimentally by NMR is on the whole correctly reproduced by the MD simulations (Figures 2 and Figure 2 – figure supplement 3). The trend for the cap loop is consistent with experiment; the S^2^ values for the cap-loop region for both oxidized and reduced nDsbD are on par with the rest of the well-ordered protein core (Figure 2 – figure supplement 3 and source data 2). The observed cap-loop opening in the MD simulations of nDsbD_ox_ is too rare an event to lower the S^2^ values in a significant way. Nonetheless, the lowered experimental {^1^H}-^15^N NOE values for nDsbD_ox_, and the need to fit them using a model that includes a τ_e_ parameter, are consistent with the observation of loop opening events in the MD simulations for nDsbD_ox_. The good level of agreement between experimental NMR parameters and values calculated from the MD trajectories provides confidence that the 1 μs simulations of oxidized and reduced nDsbD allow us to glean a realistic picture of the fast timescale dynamics of nDsbD with atomic resolution.

### NMR and MD simulations reveal local frustration in nDsbD_*ox*_

The differences in the cap loop dynamics, with the loop being more flexible in nDsbD_ox_ than in nDsbD_red_, must be a direct effect of the oxidation state of the two cysteine residues in the active site of nDsbD. X-ray crystallography (Mavridou et al. 2011) and solution NMR show that the average structures of the two redox states are very similar, with the only major difference between the two states being the presence of a disulfide bond in the active site of nDsbD_ox_ in lieu of a pair of thiol groups in nDsbD_red_.

Analysis of the MD simulations shows that the disulfide bond introduces local structural frustration in nDsbD_ox_. The side chains of C103 and C109, which form the disulfide bond, adopt gauche-conformations (χ_1_ ∼ -60°) in X-ray structures. This conformation is maintained throughout the MD simulations with only rare excursions (Figure 4C). The C103-C109 disulfide bond provides a flat binding surface for the side chain of F70. This aromatic ring packs onto the disulfide bond with its sidechain in either a gauche- or a trans orientation (Figure 4A and B). These two conformations are populated approximately equally and exchange rapidly in MD, on a timescale of tens of ns (Figure 4 – figure supplement 1). This conformational averaging is evidenced experimentally by the almost identical F70 Hβ chemical shifts, which differ by only 0.015 ppm. TALOS-N (Shen and Bax 2013) analysis of nDsbD_ox_ chemical shifts does not predict a single fixed χ_1_ value for F70, consistent with the averaging between a gauche- and a trans orientation seen for this side chain in MD simulations.

**Figure 4.**
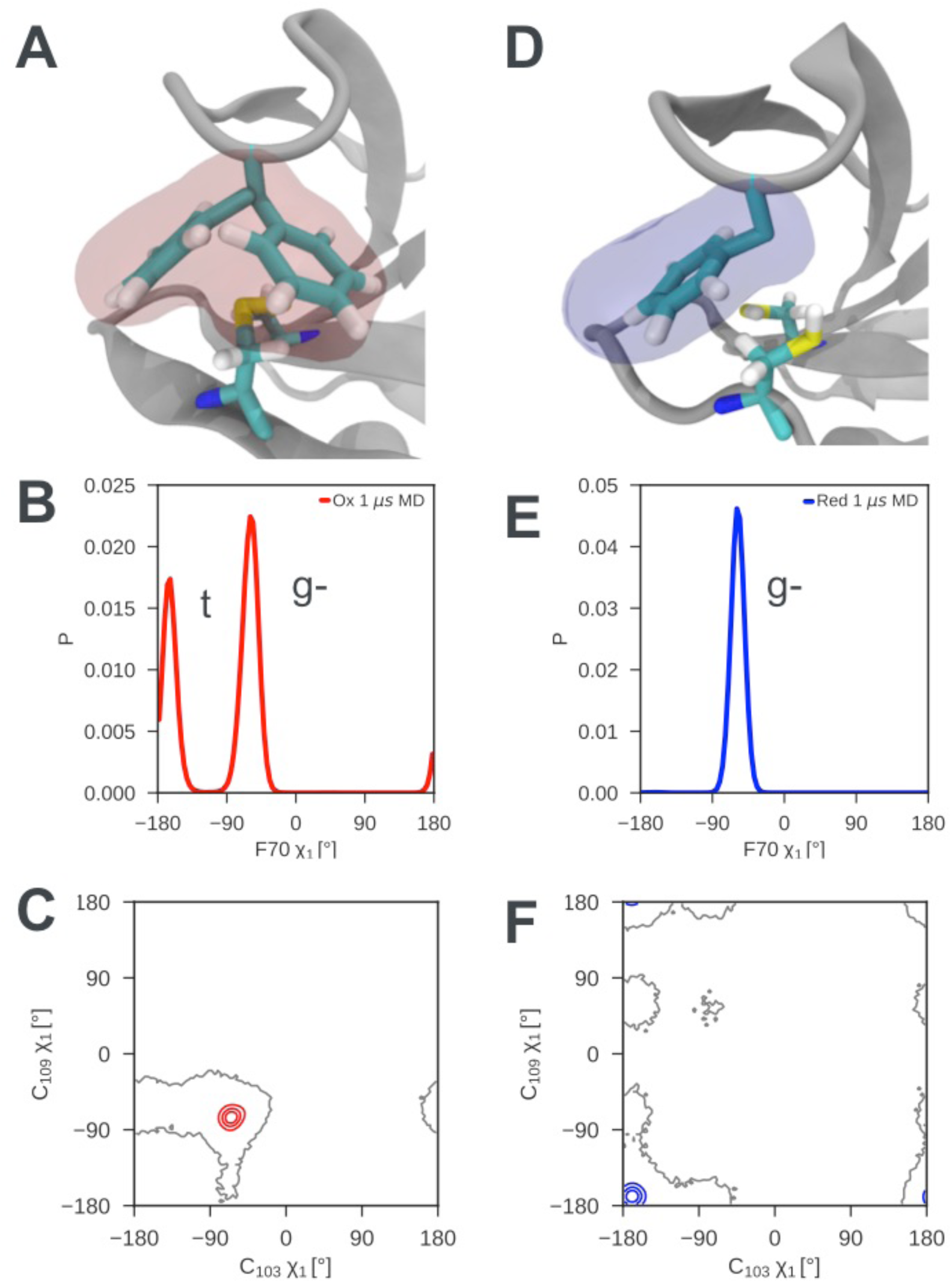
Local frustration destabilizes the cap loop of oxidized nDsbD while the cap loop of reduced nDsbD packs stably onto the active site. (A) In nDsbD_ox_, the side chain of F70 switches between a gauche- and trans orientation in MD simulations as shown by the mass density calculated for F70 (red). (B) The gauche- and trans conformations are approximately equally populated in the 1 μs MD simulations of nDsbD_ox_. (D) The cap loop and the active site of nDsbD_red_ are rigid as shown by the mass density calculated for F70 (blue). The well-defined density closely circumscribes the position of the F70 side chain in the 3PFU crystal structure. (E) F70 is locked in the gauche-conformation. The surfaces in (A) and (D) surround points with at least 0.12 relative occupancy; these structures were rendered using VMD (Humphrey and Dalke 1996). The side chains of C103 and C109 in the active site of nDsbD adopt different conformations in the oxidized (C) and reduced (F) isoforms as shown by contour plots of their χ_1_ dihedral angles. Here 1 μs of MD simulations is analysed for each redox state.

By contrast, in nDsbD_red_ F70 packs onto the active site in a single well-defined conformation, explaining why the cap loop is rigid in nDsbD_red_. The side chains of C103 and C109 both adopt trans conformations (χ_1_ ∼180°) in the X-ray structure of nDsbD_red_. This conformation is maintained throughout the MD simulations of nDsbD_red_ with only rare excursions (Figure 4F). This trans conformation of the cysteines, along with the fact that the two thiols are bulkier than a disulfide bond, result in the cysteine thiols providing a snug-fitting binding site for F70 (Figure 4D) in which one of the C109 Hβ interacts closely with the aromatic ring of F70. Consequently, F70 is locked in its gauche-conformation (Figure 4E and Figure 4 – figure supplement 1). NMR data support this packing arrangement of the cap loop onto the active site of nDsbD_red_. The Hβ of C109 are upfield shifted at 1.83 and 0.66 ppm, from a random coil value of ∼3 ppm, consistent with the close proximity to the aromatic ring of F70. The two distinct Hβ shifts for F70 of nDsbD_red_, which are separated by ∼0.2 ppm, show NOEs of differing intensity from the backbone H^N^, consistent with a single conformation for the F70 side chain. TALOS-N (Shen and Bax 2013) analysis of chemical shifts predicts a gauche-χ_1_ conformation for F70, in agreement with the MD simulations.

Recently determined X-ray structures of nDsbD from *Neisseria meningitidis* (Smith et al. 2018) provide support for our observation of oxidation-state-dependent differences in the dynamics of the cap loop. Despite sharing only 27% sequence identity with the *E. coli* domain, nDsbD_red_ from *N. meningitidis* shows a very similar structure with a closed “Phen cap” loop in which the aromatic ring of F66 adopts a gauche-conformation packing tightly against C106. Interestingly, nDsbD_ox_ from *N. meningitidis* crystallizes in the P2_1_3 space group with six molecules in the asymmetric unit. One of these molecules is observed to have an open “Phen cap” loop which superimposes very well with the open conformation observed in our MD simulations (Figure 4 – figure supplement 2A). The five molecules with a closed “Phen cap” loop show F66 packing onto the C100-C106 disulfide bond in both trans and gauche-orientations reminiscent of the two almost equally populated closed structures observed for *E. coli* nDsbD_ox_ in our MD simulations (Figure 4 – figure supplement 2B/C). The range of “Phen cap” loop conformations observed in *N. meningitidis* nDsbD_ox_ are likely to reflect conformations sampled by the protein in solution. Thus, local structural frustration appears to be a conserved feature for nDsbD_ox_ across many Gram-negative bacterial species.

### Structural features of cap loop opening

To understand how local structural frustration gives rise to cap loop opening in nDsbD_ox_ we characterized loop opening events in our MD simulations. First, we investigated whether cap loop opening is linked directly to one of the two frustrated F70 χ1 conformers. The opening of nDsbD_ox_ at 82.5 ns in Figure 3F coincides with a change of the F70 χ1 from gauche-(−60°) to trans (−180°) (Figures 5 and Figure 5 – figure supplement 1, 2, 3). During the 1 μs trajectory the cap loop experiences several shorter opening events; twenty events with a duration of at least 1 ns and a F70-to-C109 distance of at least 10 Å are observed (Figure 3, Figure 3 – figure supplement 2, Figure 5 – figure supplement 1 and 4). In only one of these events does F70 undergo a similar gauche- to trans transition at the point of opening. Fourteen of the openings start from F70 in the trans conformer, while five openings start from the gauche-conformer. Thus, loop opening is not underpinned by a specific F70 χ1 conformer.

**Figure 5.**
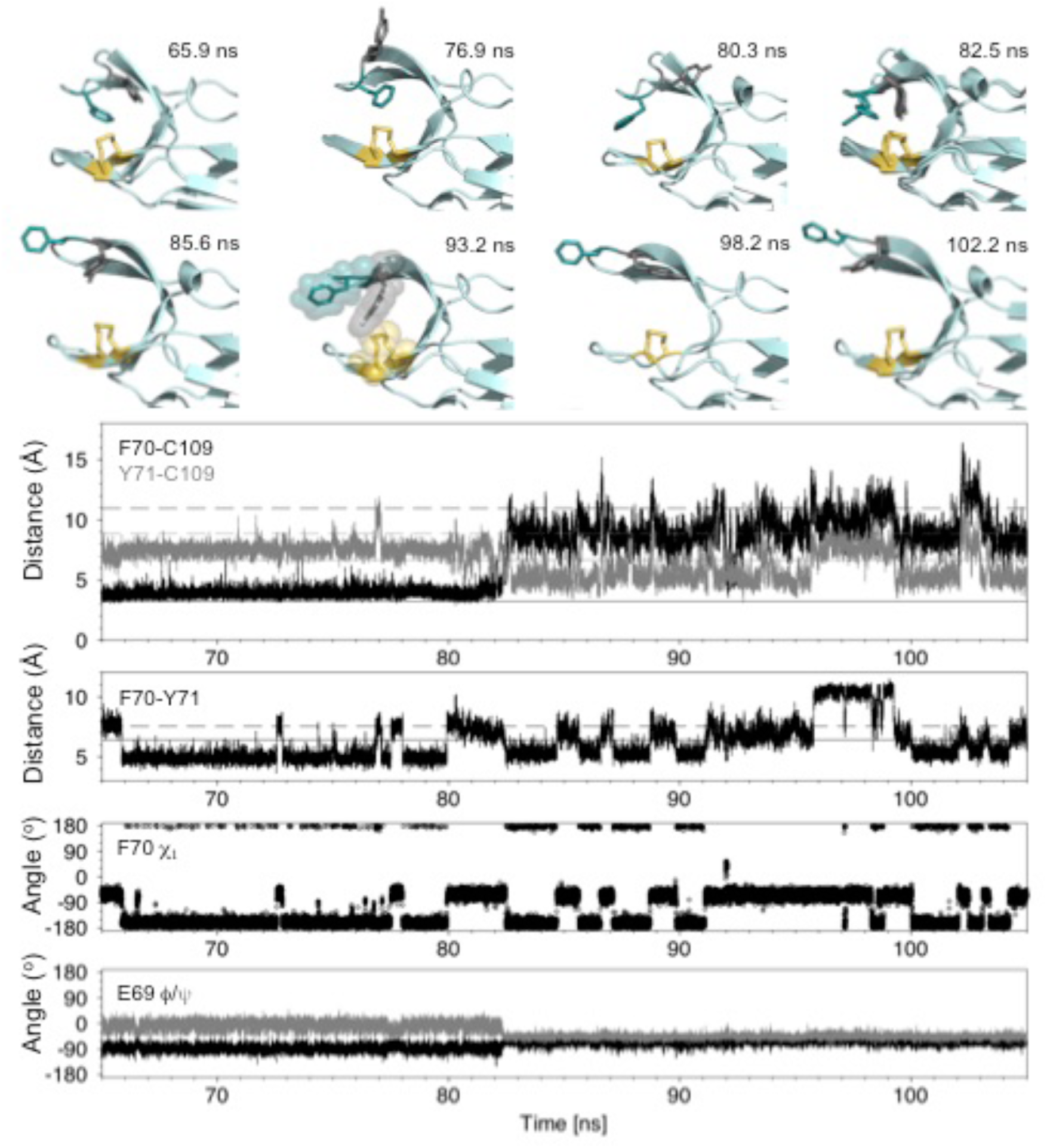
Analysis of the major cap-loop opening event observed in MD simulations of nDsbD_ox_. Top: Snapshots of structures generated from the MD simulation between 65 and 105 ns show the range of cap loop conformations and, in particular, the F70 and Y71 side chain positions that are sampled. The backbone of nDsbD is shown as a cartoon in pale cyan. F70, Y71 and C103/C109 are shown in a stick representation and coloured deep teal, grey and yellow, respectively. Two structures are overlaid in the 82.5 ns panel; these illustrate the conversion of F70 from the gauche-to the trans conformation observed simultaneously with loop opening. A surface representation is used in the 93.2 ns snapshot to illustrate how Y71 blocks closing of the cap loop. Bottom: Structural parameters are plotted as a function of simulation time. The distance from the centre of the rings of F70 and Y71 to C109 Sγ are shown in black and grey in the first panel. The distance between the ring centres of F70 and Y71 are shown in the second panel. The solid and dashed lines show these distances in the closed (1L6P) and open (1VRS) X-ray structures, respectively. In the third and fourth panels the F70 χ_1_ torsion angle and the E69 ϕ and Ψ torsion angles are shown. These and additional structural parameters are shown in the figure supplements for the entire loop opening period (60-200 ns) and for a second 22 ns loop opening event.

By contrast, we found that the dynamics of Y71 and the relative orientation of F70 and Y71 seem to be important for cap loop opening. In closed nDsbD_ox_ Y71 also samples both gauche- and trans χ1 conformers, showing a strong preference for gauche- (86%). Despite this preferred conformation, we note that there is no correlation between the χ1 values of F70 and Y71; the less common Y71 trans conformer is observed for both frustrated F70 conformers. We observed the sequence of events leading to loop opening (Figure 5 and Figure 5 – figure supplement 1,2,3). Although the cap loop remains closed from 60-80 ns, the side chains of both F70 and Y71 sample a number of different conformations. From 60 to 65 ns, both F70 and Y71 are in the gauche- χ1 conformation observed in the 1L6P X-ray structure. At 65.9 ns, F70 χ1 switches to the trans conformer. This brings the F70 and Y71 rings closer together. From 66 to 80 ns, F70 remains primarily in the trans conformer with only brief excursions to gauche- (at ∼72.7 and ∼77.7 ns), and at 79.9 ns F70 returns to the gauche- χ1 conformer. Meanwhile, at 76.9 and 80.3 ns the ring of Y71 transiently moves into more exposed conformations as a result of changes to χ1 and χ2, but the loop remains closed (Figure 5 and Figure 5 – figure supplement 2). Immediately before loop opening the F70 and Y71 rings are in close proximity, while Y71 is also close to C109. This flexible and dynamic nature of the side chains of F70 and Y71 in the closed state is reflected in their RMSD, to the equilibrated X-ray structure, observed during the MD trajectories. In the nDsbD_ox_ 1 μs trajectories, F70 and Y71 have RMSDs of 2.8 ± 0.8 and 2.6 ± 0.7 Å, respectively when the loop is closed (F70-C109 < 6 Å). By contrast, these RMSD values are 1.7 ± 0.6 and 1.6 ± 0.8 for F70 and Y71, respectively in nDsbD_red_ trajectories. It is worth noting here that the aromatic ring of F67 in *N. meningitidis* nDsbD_ox_, which is homologous to Y71 in *E. coli* nDsbD, is also dynamic, adopting different conformations in the crystallized closed structures (Figure 4 – figure supplement 2) (Smith et al. 2018). Taking these observations together, we propose that the existence of local structural frustration for F70 results in increased dynamics for both F70 and Y71, which leads to occasional loop opening in nDsbD_ox_.

Following opening of the cap loop at 82.5 ns, Y70 and Y71 continue to sample a variety of conformations, which could be important for the cap loop to access a fully open structure. The initial open states show the ring of F70 pointing inwards, due to its trans conformation. Transition of F70 to the gauche-conformer after loop opening, leads to a more extended orientation for F70. The extent of loop opening, as measured by the F70-to-C109 distance, fluctuates during the 120 ns open period; for example, at 85.6 ns the loop is in a more open conformation (12.5 Å) than at 93.2 ns (8.6 Å) (Figure 5). Interestingly, as the F70-to-C109 distance decreases, the loop is often prevented from closing fully by the side chain of Y71 which sits in between C109 and F70 in a ‘blocking’ conformation (for example at 93.2 ns and later at 151.9 and 155.8 ns (Figure 5 and Figure 5 – figure supplement 5)). The orientation of F70 relative to Y71 also changes while the loop is open. At 98.2 ns the position of Y70 is similar to that observed in the open 1VRS X-ray structure but Y71 points inwards. At 102.2 ns the Y70 and F71 orientations are more similar to the 1VRS structure (Figure 5). The cap loop can also adopt conformations that are more open than the open 1VRS structure (at ∼145 and ∼181 ns); these structures are characterized by larger F70-C109 and Y71-C109 distances (> 16 Å and 10 Å, respectively) (Figure 5 – figure supplement 1 and 5). These conformations may be important for the initial docking of cDsbD prior to nDsbD/cDsbD complex formation, as they would minimize the steric clashes between the two domains during binding.

The backbone ϕ and Ψ torsion angles of D68, E69, F70, Y71 and G72 also show clear changes at 82.5 ns when the loop begins to open (Figure 5 and Figure 5 – figure supplement 3). E69 ϕ and Ψ become less variable in the open loop while the ϕ and Ψ values for D68 and F70 become more variable. These torsion angle changes are correlated with loss of the hydrogen bond between the amide nitrogen of Y71 and the side chain carboxyl group of D68 upon loop opening. This hydrogen bond is observed in the 1L6P X-ray structure and in 95% of structures between 60 and 82.5 ns, prior to loop opening. Notably, D68 and E69 adopt ϕ and Ψ values more reminiscent of the closed state between 160 and 180 ns, despite the loop remaining open (Figure 5 – figure supplement 3); during this period the Y71HN-D68Oδ hydrogen bond is again observed in 62% of structures. By contrast, the backbone hydrogen bonds which define the antiparallel β-hairpin of the cap loop and which involve D68-G72 and H66-S74, are always retained in the open loop structures.

We have also investigated the dynamics of Y71 in the simulations of loop closing, and in particular those simulations in which the loop fails to close completely within 10 ns. During these trajectories Y71 adopts conformations in which it sits between F70 and C109 thus blocking complete closure of the loop as observed, for example, at 93.2 ns in the major opening event and at 2.8 ns in the incomplete loop closing seen in Figure 3G (Figure 5 and Figure 5 – figure supplement 5).

In the light of the important roles identified for both F70 and Y71 in cap loop opening, we investigated the conservation of cap loop residues using bioinformatics. We have used 134 representative sequences, one from each bacterial genus, for our sequence alignment (Figure 5 – figure supplement 6). We find that the cap-loop motif is more conserved than the flanking β-strands, in agreement with the central role that this structural motif plays in nDsbD function. In 53% of the sequences an F or a Y is found at position 70; in a further 36.4% of the sequences a residue with a relatively bulky side chain (K/I/L/E/V/T/N), which would shield the cysteines from solvent, is found at position 70. Interestingly, in 85% of sequences an F or a Y is found at position 71; in a further 5.2% of the sequences position 71 is occupied by a bulky hydrophobic residue (I or L). This high conservation of an aromatic amino acid observed for this position further highlights the key role played by Y71 in contributing to loop opening and, more importantly, in preventing rapid loop closure, thus likely facilitating the interaction of nDsbD with cDsbD prior to reductant transfer.

### NMR relaxation dispersion experiments

Using ^15^N NMR relaxation dispersion experiments (Baldwin and Kay 2009, Loria, Rance, and Palmer 1999) we also detected a difference in the μs-ms timescale dynamics of reduced and oxidized nDsbD. For nDsbD_red_, no evidence for μs-ms dynamics in the cap loop or the active site were found (Figure 6A-C). This observation is consistent with the absence of a significantly populated open state of nDsbD_red_. By contrast, in nDsbD_ox_, eight residues, V64, W65, E69, F70, Y71, G72, K73 and S74, all in the cap-loop region, showed strong relaxation dispersion effects (Figure 6D-F and 7A). These results are consistent with previous observations about peak intensities in ^1^H-^15^N HSQC spectra (Mavridou et al. 2012). The peaks of C103, C109 and Y110 are not visible and the peak of A104 is very weak in the spectrum of nDsbD_ox_, but not nDsbD_red_ (Mavridou et al. 2012), likely due to extensive chemical exchange processes in nDsbD_ox_ leading to ^1^H^N^ and/or ^15^N peak broadening. Thus, the cap loop of nDsbD_ox_, and likely also the active-site cysteines, exchange between different conformations on the μs-ms timescale, whereas nDsbD_red_ adopts a single conformation on this timescale.

**Figure 6.**
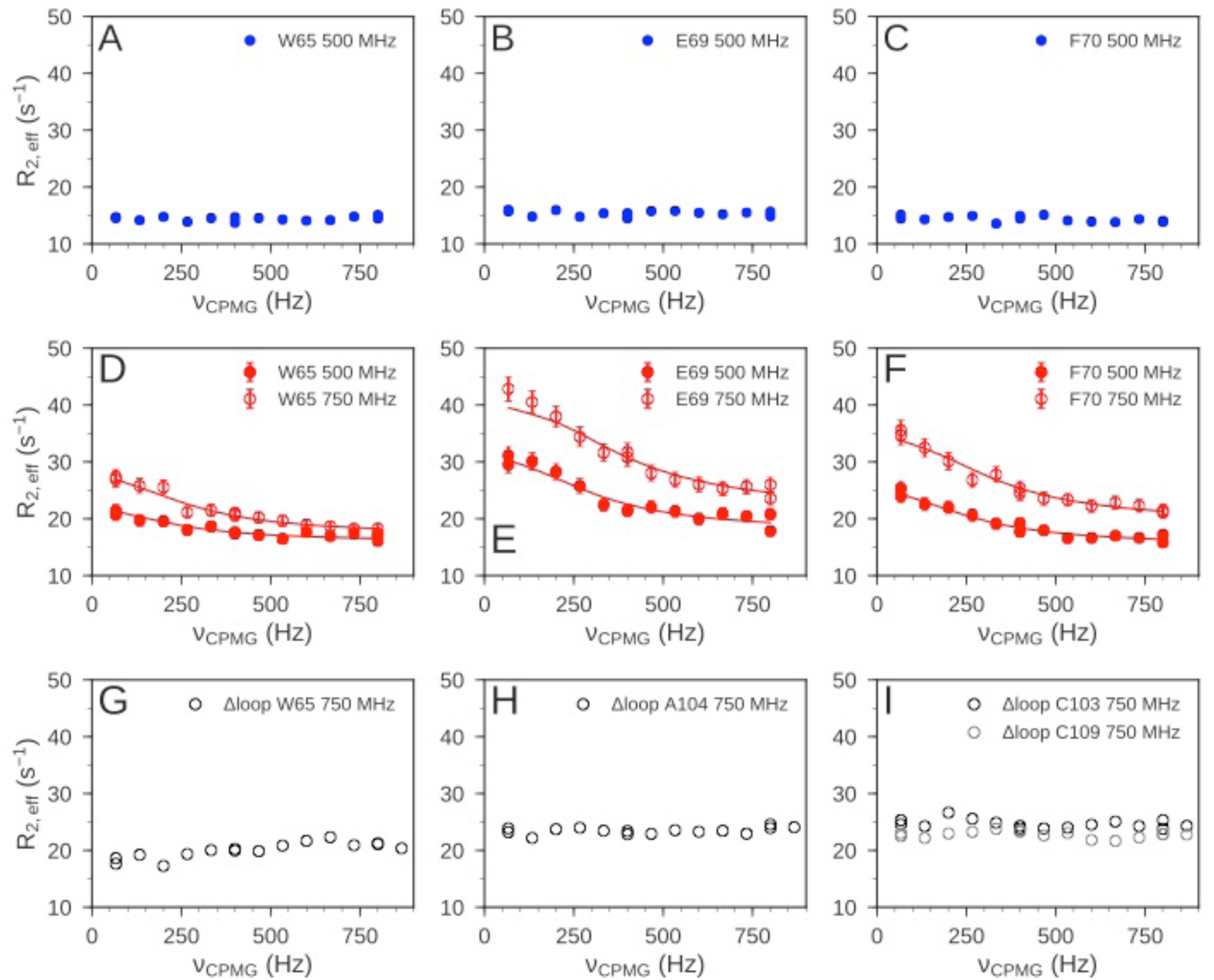
^15^N relaxation dispersion experiments were used to determine the μs-ms dynamics of nDsbD. For nDsbD_red_, flat relaxation dispersion profiles were obtained as illustrated for (A) W65, (B) E69 and (C) F70 in the cap loop. For nDsbD_ox_ clear relaxation dispersion effects were measured for (D) W65, (E) E69 and (F) F70; error bars were determined as described in source data 1. In the oxidized cap-loop deletion mutant (Δloop-nDsbD_ox_), flat relaxation dispersion profiles were obtained for (G) W65, (H) A104 and (I) C103/C109 in the active site. Experimental data are represented by circles and fits to the data for wild-type nDsbD_ox_ are shown by solid lines.

Analysis of the relaxation dispersion data collected at 25°C (using CATIA (Vallurupalli, Hansen, and Kay 2008, Baldwin, Hilton, et al. 2011, Alderson et al. 2019)) showed that nDsbD_ox_ undergoes a single global exchange process, between a dominant major state (p_A_ 98%) and an alternative minor state (p_B_ 2%) (Figure 6 – source data 1). To corroborate that a single exchange process gives rise to the relaxation dispersion curves, we repeated the experiments at four additional temperatures (10, 15, 20 and 35 °C). If the residues underwent different exchange processes we would expect them to show a differential response to temperature (Grey, Wang, and Palmer 2003). The relaxation dispersion data for all residues in the cap loop recorded at the five temperatures could be fitted simultaneously (Baldwin, Hilton, et al. 2011, Alderson et al. 2019) (Figure 6 – figure supplement 1A and source data 1). This confirmed that a single process underlies the μs-ms chemical exchange in the cap loop of nDsbD_ox_. In addition, the analysis of the data at all five temperatures yielded a detailed description of the thermodynamics and kinetics of the exchange process (Figure 6 – figure supplement 1B and source data 1). The minor “excited” state is enthalpically, but not entropically, more favorable than the major ground state of nDsbD_ox_. The fits also demonstrated that no enthalpic barrier separates the ground and excited state of nDsbD_ox_. This absence of an enthalpic barrier between the two states might mean that no favorable interactions have to be broken in the transition state, but that the diffusion process itself may limit the speed of the transition (Dill and Bromberg 2010).

Analysis of the chemical shift differences determined from the relaxation dispersion experiments suggests that the conformation adopted by the cap loop in the minor state of nDsbD_ox_ might be similar to the conformation of nDsbD_red_. To interpret the chemical shift differences we calculated the shift changes expected if the minor state features a disordered cap loop (Nielsen and Mulder 2018). Such a state would include more open conformations and such structures might facilitate the binding of the nDsbD interaction partners. The lack of correlation in Figure 7B demonstrates that the cap loop does not become disordered in the less populated minor state. The ^15^N chemical shift differences between the major and minor states do, however, resemble the chemical shift differences between the oxidized and reduced isoforms of nDsbD (Figure 7C) (Mavridou et al. 2012). Nonetheless, the correlation in Figure 7C is not perfect; the differences between the chemical shifts of the minor state of nDsbD_ox_ and those of nDsbD_red_ likely reflect the local electronic and steric differences between having a disulfide bond (in the minor state of nDsbD_ox_) and two cysteine thiols (in nDsbD_red_) in the active site. Importantly, the sign of the chemical shift differences between the major and minor state, which could be determined experimentally for W65, E69, F70, Y71, G72 and K73, further indicated that the minor state structure of the cap loop of nDsbD_ox_ is not disordered (Figure 7D and Figure 7 – source data 1), but instead may be similar to that adopted in nDsbD_red_ (Figure 7E and Figure 7 – source data 1).

**Figure 7.**
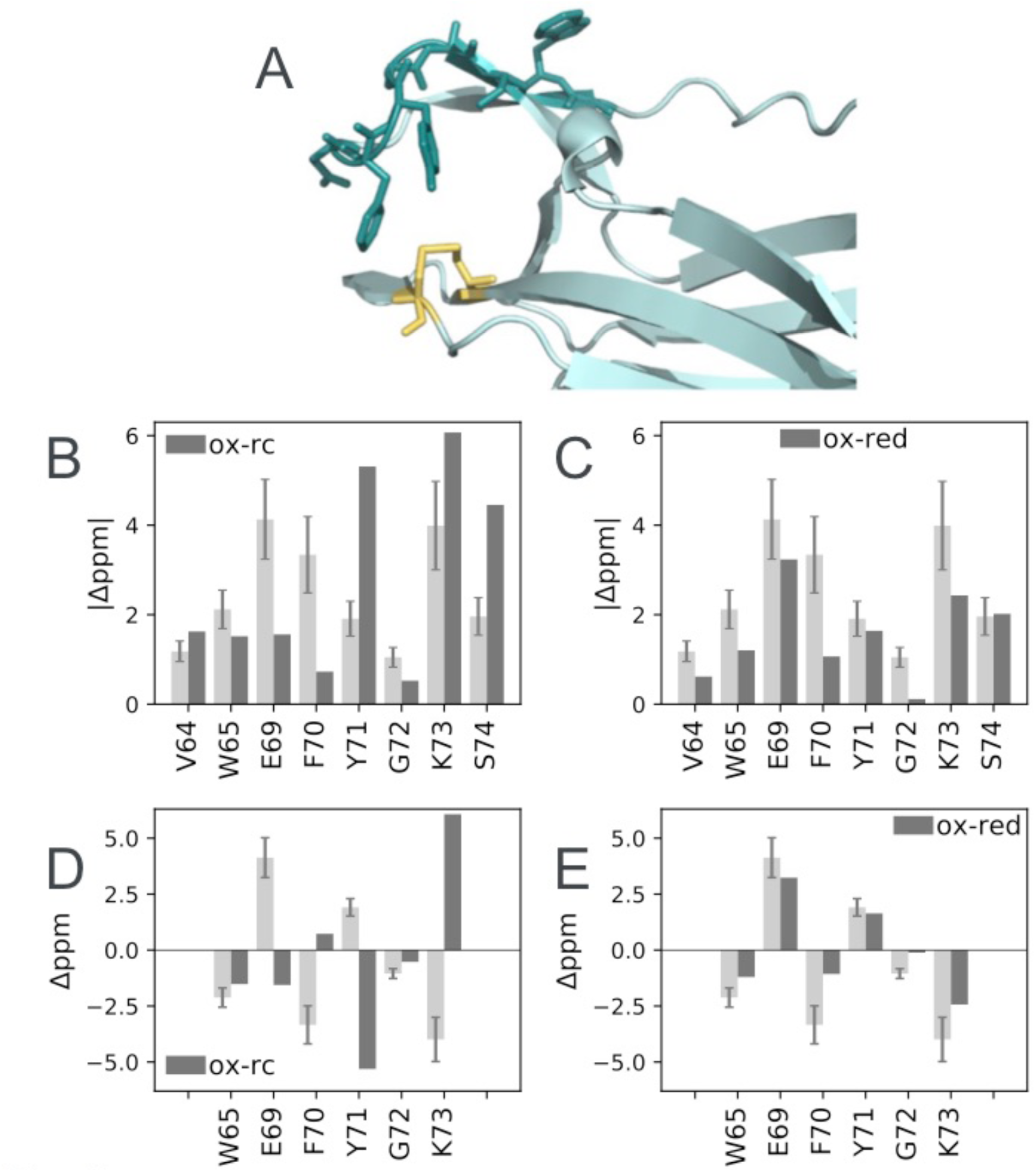
Structural characterization of μs-ms dynamics in nDsbD_ox_. (A) Residues 64, 65 and 69-74 with chemical shift differences between the major and minor states of > 1 ppm are shown in a stick representation in dark teal. C103 and C109 are shown in yellow. The magnitude of the ^15^N chemical shift differences between the major and minor states are compared in (B) with the magnitude of predicted shift differences between the native and random coil states and in (C) with the magnitude of the experimental shift difference between nDsbD_ox_ and nDsbD_red_. Shift differences between the major and minor states for W65, E69, F70, Y71, G72 and K73, for which the sign has been obtained experimentally, are compared with the predicted shift differences between the native and random coil states in (D) and between nDsbD_ox_ and nDsbD_red_ in (E). The experimentally determined chemical shift differences are shown in light grey, with error bars. Random coil shifts were predicted using POTENCI (Nielsen and Mulder 2018). Details of the fitting procedures used for the dispersion data and the determination of errors in the chemical shift differences are presented in Figure 6 – figure supplement 1 and source data 1 and in Figure 7-source data 1.

### Coupling between the active site and the cap loop gives rise to the alternative minor state of oxidized nDsbD

The alternative minor state of nDsbD_ox_, which we detected by NMR relaxation dispersion, can also be attributed to coupling between the dynamics of the active-site cysteine residues and the cap loop. In this alternative minor state, the C103 and C109 side chains likely adopt a ‘reduced-like’ conformation. Both side chains adopt a gauche-conformation (χ_1_ ∼ -60°) in nDsbD_ox_, but a trans conformation (χ_1_ ∼ 180°) in nDsbD_red_. In MD simulations of both redox states, rare excursions of one or the other χ_1_, but never both, to the alternative conformation are observed (Figure 4 C/F). We postulate that in the minor state of nDsbD_ox_, both of the cysteine side chains adopt the ‘reduced-like’ trans conformation. In this state, the cap loop could pack tightly onto the active-site disulfide bond, as observed in MD simulations of nDsbD_red_. The backbone ^15^N chemical shifts of this minor state consequently resemble those of the reduced protein. Model building of a disulfide bond into the reduced 3PFU crystal structure followed by energy minimization with XPLOR (Brunger 1992) showed that the resulting ‘oxidized’ structure maintains the ‘reduced-like’ trans conformation (χ_1_ ∼ -140°) for the cysteine side chains and the gauche-conformation for F70 (Figure 8B). This ‘reduced-like’ conformation for nDsbD_ox_, with a more tightly packed active site and a cap loop which lacks frustration, might be enthalpically more favorable and entropically less favorable than the major state of nDsbD_ox_ as shown by the analysis of relaxation dispersion data collected at multiple temperatures (Figure 6 – figure supplement 1 and source data 1). Previously, NMR relaxation-dispersion experiments and MD simulations have detected complex dynamics in BPTI stemming from one of its disulfide bonds (Grey, Wang, and Palmer 2003, Shaw et al. 2010, Xue et al. 2012), further suggesting that the disulfide bond shapes the μs-ms dynamics of the active-site and cap-loop regions of nDsbD_ox_.

**Figure 8.**
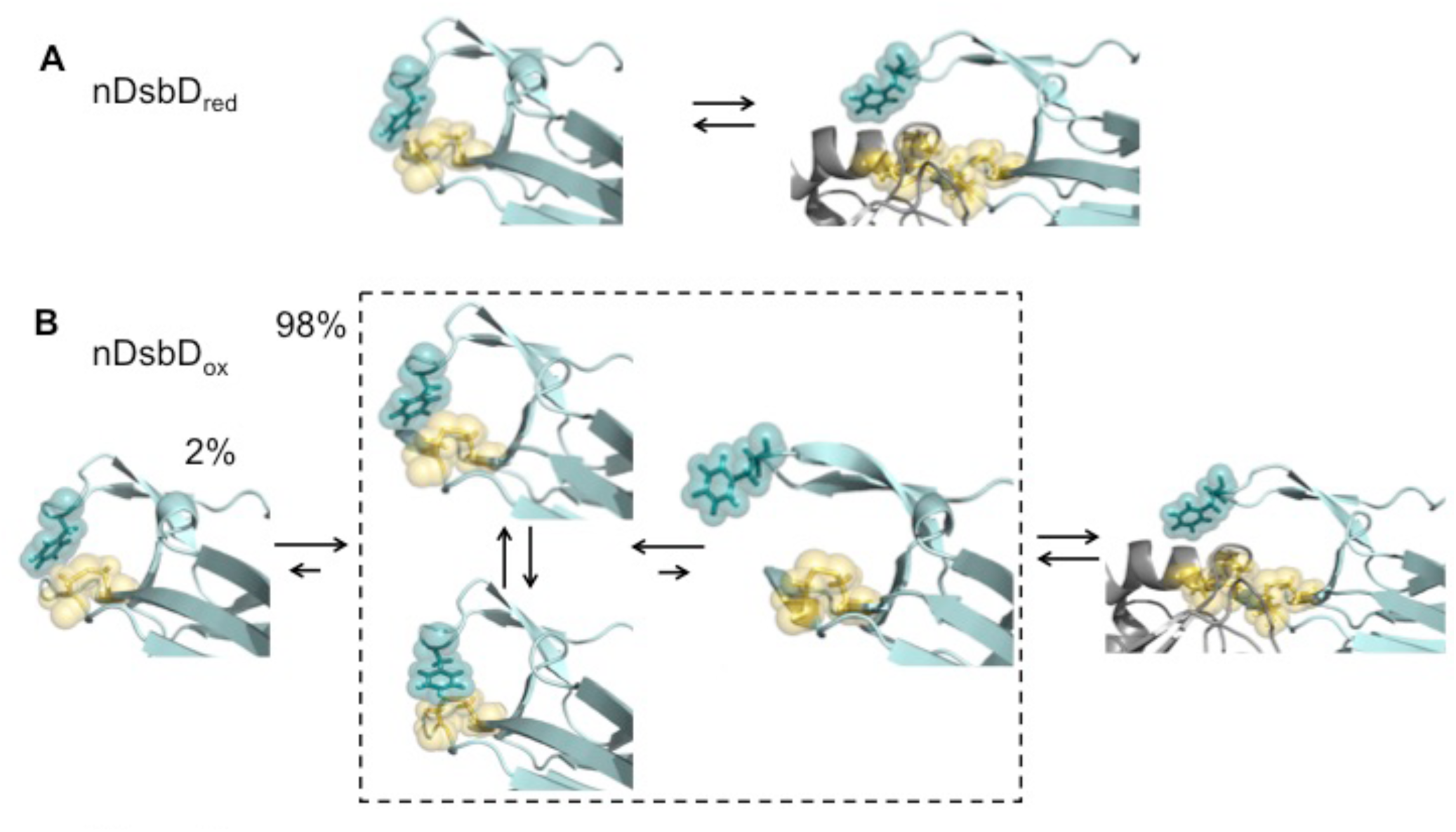
Local frustration occurs in oxidized, but not in reduced, nDsbD. (A) The cap loop in nDsbD_red_ maintains a closed conformation shielding the active-site cysteine thiols. The cap loop may only open to expose the cysteine thiols via an induced-fit mechanism when it encounters its legitimate binding partners, CcmG, DsbC and DsbG. (B) nDsbD_ox_ undergoes more complex dynamics as a result of local structural frustration. F70 interacts with C109 Sγ both in its gauche- (top left structure in dashed box) and trans (lower left structure in dashed box) orientations. These mutually competing interactions destabilize the packing of the cap loop onto the active site. Consequently, open conformations (right-hand structure in dashed box) are accessible in nDsbD_ox_ and these would allow nDsbD_ox_ to form complexes with its legitimate binding partner, cDsbD_red_, via a conformational selection mechanism. The left-hand structure shows a reduced-like minor (2%) conformation identified by relaxation dispersion experiments and populated on a slower timescale. The backbone of nDsbD and cDsbD are shown in cyan and grey, respectively. F70 (dark teal), C103 and C109 (yellow) are shown with sticks and a surface representation. (A) is from the 3PFU X-ray structure of nDsbD_red_, the left-hand panel in (B) is the energy minimized 3PFU structure with a disulfide bond incorporated, the structures in the dashed rectangle in (B) are ‘snapshots’ from the MD simulations of nDsbD_ox_. The bound complexes shown on the right in (A) and (B) are based on the 1VRS X-ray structure.

### NMR relaxation dispersion experiments using a loop deletion variant

We confirmed the tight coupling between the dynamics of the cap loop and the active site by NMR relaxation dispersion experiments using an nDsbD variant. We generated an nDsbD protein with a truncated cap loop in which the eight residues H66-K73 were replaced by A-G-G (Δloop-nDsbD). The X-ray structure of oxidized Δloop-nDsbD shows that deletion of the cap loop does not cause any significant changes in structure; a Cα RMSD of 0.25 Å is obtained for residues 12-122 (Figure 6 – figure supplement 2 and source data 2). Nevertheless, in the absence of F70, the shorter cap loop can no longer closely pack onto the disulfide bond. As a result, no relaxation dispersion is detected in the truncated cap loop variant, as shown for W65 (Figure 6G). For A104 (A99), a much sharper amide peak is observed in the HSQC spectrum of Δloop-nDsbD_ox_ and this residue shows a flat dispersion profile (Figure 6H). Importantly, the ^1^H^N^-^15^N peaks for C103 (C98) and C109 (C104), which are not observed in the wild-type oxidized protein, can be detected for Δloop-nDsbD_ox_. Finally, ^15^N relaxation dispersion profiles for these cysteines are also flat (Figure 6I). Thus, truncation of the cap loop alters the μs-ms dynamics of the cap-loop region and the active-site cysteines, further supporting our proposed model of tight coupling between the cap loop and active-site cysteines.

## Conclusion

We have studied oxidized and reduced nDsbD using NMR experiments and microsecond duration MD simulations, and have demonstrated that the conformational dynamics of the active-site cysteines and the cap loop are coupled and behave in an oxidation-state-dependent manner. We showed that the cap loop is rigid in nDsbD_red_, whilst it exhibits more complex dynamics in nDsbD_ox_ (Figure 8). Although the cap loop is predominately closed and ordered in both oxidation states, for nDsbD_red_ we found no evidence for a significant population with an open cap loop. Our results argue that the cap loop of nDsbD_red_ remains closed at all times, to protect the active-site cysteine thiol groups from non-cognate reactions in the oxidizing environment of the periplasm. As such, the nDsbD_red_ cap loop may only open to expose the cysteine thiols via an induced-fit mechanism when it encounters its legitimate binding partners, for example CcmG, DsbC and DsbG. In nDsbD_ox_, on the other hand, the cap loop opens spontaneously in MD simulations and while fully open conformations of the cap loop tended to close, partially open structures were stable. Thus, nDsbD_ox_ may form a complex with its sole legitimate binding partner, cDsbD_red_, via a conformational selection mechanism. Notably, NMR relaxation dispersion experiments provided no evidence for a long-lived open or disordered conformation of the cap loop as might have been expected. Instead, a minor state with a ‘reduced-like’ conformation in the active site and cap loop is observed solely for nDsbD_ox_.

Importantly, combining NMR experiments and MD simulations, we showed how local frustration (Ferreiro et al. 2007, 2011) in nDsbD_ox_, but not in nDsbD_red_, gives rise to differences in the dynamics of the two redox isoforms, on multiple timescales, providing insight into how this could affect protein function. Local frustration has been highlighted in the interaction interfaces of many protein complexes (Ferreiro et al. 2007), where mutually exclusive favorable interactions result in a dynamic equilibrium of competing structures and determine which parts of a protein are flexible. In nDsbD_red_, the cap loop packs perfectly onto the active-site cysteines and the closed conformation is stable, something that is crucial in the very oxidizing environment in which the domain has to provide electrons. By contrast, in nDsbD_ox_, the disulfide bond disrupts this tight fit of the aromatic ring of F70. Consequently F70 is no longer locked in a single conformation and switches freely between gauche- and trans orientations. This results in increased flexibility for both F70 and Y71 and drives occasional opening of the cap loop, which in this oxidation state might facilitate nDsbD_ox_ receiving electrons from its partner. This is further enabled by a possible role of Y71 in preventing rapid loop closing. The observation from sequence alignments that the general architecture of the cap-loop region in β- and γ-proteobacteria is conserved, highlights that our observations in *E. coli* nDsbD can be generalized for many other bacterial species where reductant provision is driven by DsbD in the cell envelope.

Overall, our results show that it is the oxidation state of the cysteine pair in nDsbD that acts as a ‘switch’ controlling the presence or absence of local frustration in its active site. This oxidation-state-dependent ‘switch’ determines the dynamic behavior of the cap loop which translates into optimized protein-protein interaction and biological activity. We propose that redox-state modulation of local frustration, such as that observed here for nDsbD, may play a wider role in biomolecular recognition in other pathways involving redox proteins which are ubiquitous in all organisms. Ultimately, the understanding gained from studies like this would allow controlled introduction of frustration, mimicking intricate natural systems such as DsbD, for more translational applications such as the production of synthetic molecular devices.

## Materials and Methods

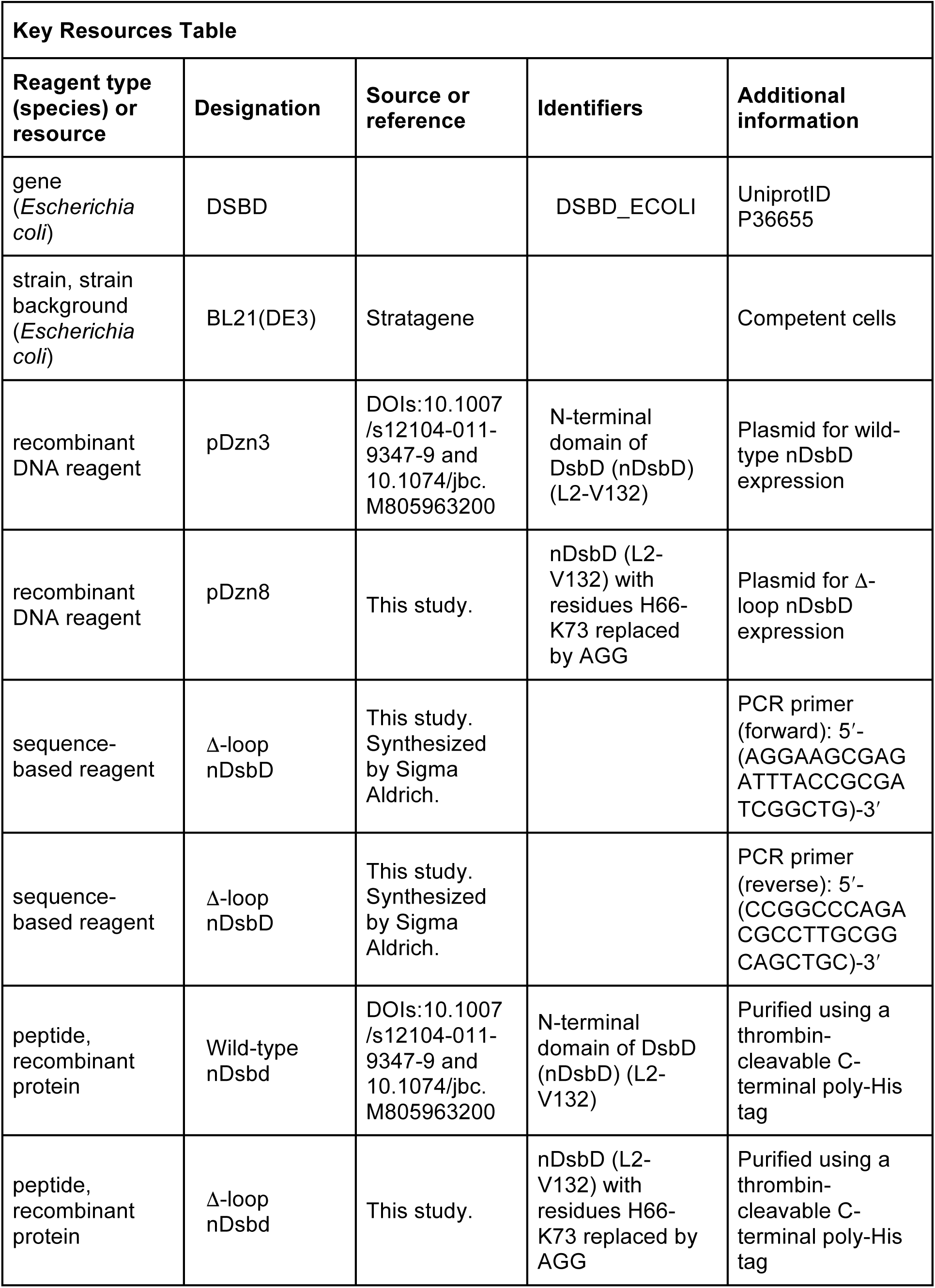
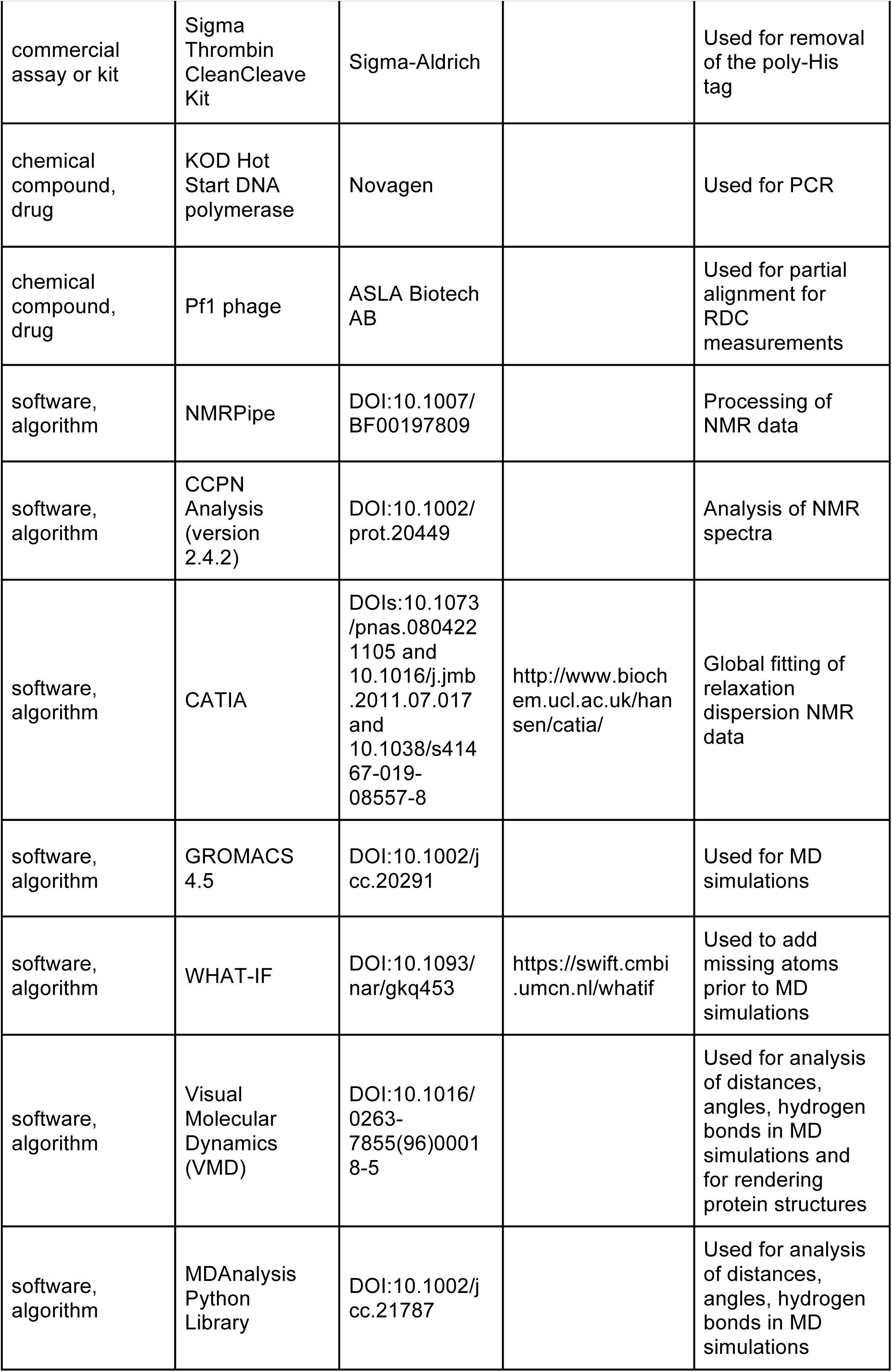
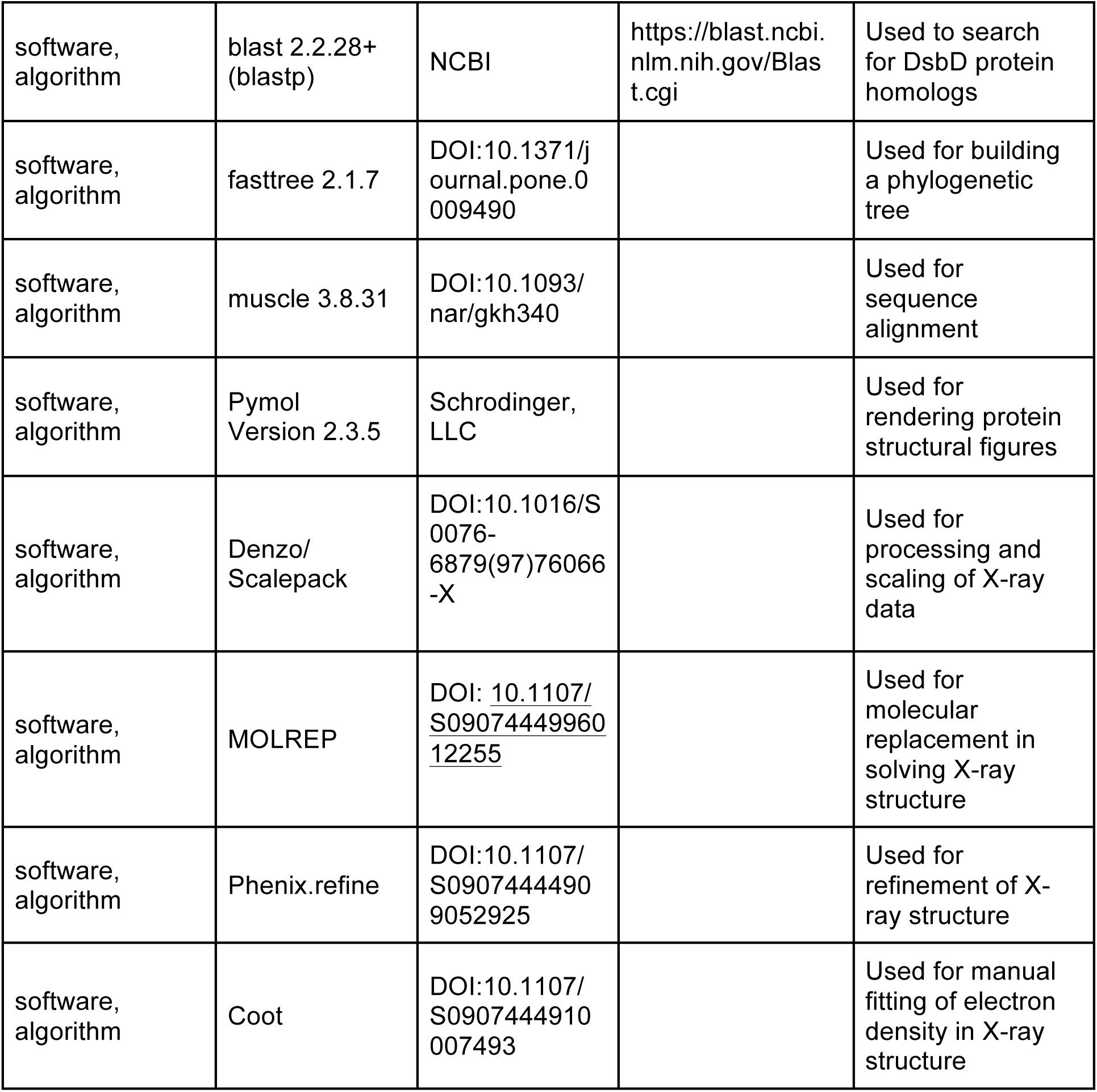

### Construction of plasmids and protein production

The plasmid pDzn3, described in previous work (Mavridou et al. 2012, Mavridou et al. 2009), encodes isolated wild-type nDsbD (L2-V132) bearing a thrombin-cleavable C-terminal poly-histidine tag. This construct was used as a template to produce a cap-loop deletion variant of nDsbD (Δloop-nDsbD), where residues H66-K73 were replaced by the amino acid sequence A-G-G. Site-directed mutagenesis (ExSite, Stratagene) was performed using oligonucleotides 5’-(AGGAAGCGAGATTTACCGCGATCGGCTG)-3’ and 5’-(CCGGCCCAGACGCCTTGCGGCAGCTGC)-3’. PCR was performed with KOD Hot Start DNA polymerase (Novagen) and oligonucleotides were synthesized by Sigma Aldrich. DNA manipulations were conducted using standard methods. The resulting plasmid was named pDzn8. nDsbD and Δloop-nDsbD were expressed using BL21(DE3) cells (Stratagene) and purified from periplasmic extracts of *E. coli* using a C-terminal poly-histidine tag, as described previously (Mavridou et al. 2012, Mavridou et al. 2007). Thrombin cleavage of the affinity tag was performed using the Sigma Thrombin CleanCleave Kit (Sigma) according to the manufacturer’s instructions.

### NMR spectroscopy

NMR experiments were conducted using either home-built 500, 600 and 750 MHz spectrometers equipped with Oxford Instruments Company magnets, home-built triple-resonance pulsed-field gradient probeheads and GE/Omega data acquisition computers and software, or using a 750 MHz spectrometer equipped with a Bruker Avance III HD console and a 5mm TCI cryoprobe. All experiments were conducted at 25 °C and at pH 6.5 in 95% H_2_O/5% D_2_O, unless stated otherwise. Spectra were processed using NMRPipe (Delaglio et al. 1995) and analyzed using CCPN Analysis (Skinner et al. 2016, Vranken et al. 2005).

Residual dipolar couplings (RDCs) were measured using Pf1 phage (ASLA Biotech AB) on separate samples for oxidized and reduced nDsbD. RDCs were measured at 600 MHz for 0.5 mM nDsbD with 10 mg/ml Pf1 phage, 2 mM K_2_PO_4_, 0.4 mM MgCl_2_, 0.01% NaN_3_ and 10% D_2_O. Measurements were also carried out for isotropic solutions prior to the addition of the Pf1 phage. RDCs were measured using the InPhase-AntiPhase (IPAP) approach (Ottiger, Delaglio, and Bax 1998). The F_2_ ^1^H and F_1_ ^15^N dimensions were recorded with 128 scans per increment, sweep widths of 7518.8 and 2000.0 Hz, respectively, and 1024 and 128 complex points, respectively. The quality factor (Q) that describes the agreement between calculated and observed RDCs was determined using the procedure of Ottiger et al. (Ottiger, Tjandra, and Bax 1997).

{^1^H}-^15^N heteronuclear NOE, ^15^N T_1_ and T_2_ data (Boyd, Hommel, and Campbell 1990, Kay, Torchia, and Bax 1989, Palmer et al. 1992) were measured at 600 MHz. The {^1^H}-^15^N heteronuclear NOE was also measured at 500 MHz. For the T_1_ measurement, spectra with 14 different relaxation delays, ranging from 20 ms to 2 s, were collected. The T_2_ was measured by recording spectra with 14 relaxation delays between 8 ms and 400 ms, with a Carr-Purcell Meiboom Gill (CPMG) delay τ_CPMG_ of 0.5 ms. For the T_1_ and T_2_ measurements, a recycle delay of 2 s was used. The {^1^H}-^15^N NOE was measured by comparing peak heights in interleaved spectra recorded with and without saturation of the protons for 3 and 4 s at 500 and 600 MHz, respectively. The F_2_ ^1^H and F_1_ ^15^N dimensions were recorded with sweep widths of 7518.8 and 1785.7 Hz, respectively, and with 1024 and 128 complex points, respectively. The T_1_ and T_2_ and NOE experiments were collected with 16, 16 and 128 scans per increment, respectively.

^15^N relaxation data were collected for 107 and 117 of the 126 non-proline residues of oxidized and reduced nDsbD, respectively. Data for the remaining residues could not be measured due to weak or overlapping peaks. T_1_, T_2_ and {^1^H}-^15^N NOE values were determined as described previously (Smith et al. 2013). Uncertainties in the T_1_, T_2_ and {^1^H}-^15^N NOE values were estimated from 500 Monte Carlo simulations using the baseline noise as a measure of the error in the peak heights (Smith et al. 2013). Relaxation data were analyzed using in-house software (Smith et al. 2013); this incorporates the model-free formalism of Lipari and Szabo (Lipari and Szabo 1982a, b) using spectral density functions appropriate for axially-symmetric rotational diffusion (Abragam 1961) and non-colinearity of the N-H bond vector and the principal component of the ^15^N chemical shift tensor (Boyd and Redfield 1998) with model selection and Monte Carlo error estimation as described by Mandel et al. (Mandel, Akke, and Palmer 1995). Calculations were carried out using an N-H bond length of 1.02 Å, a ^15^N chemical shift anisotropy, (σ_‖_ - σ_⊥_), of -160 ppm, and a D_‖_/D_⊥_ ratio of 2.0.

^15^N relaxation-dispersion experiments (Loria, Rance, and Palmer 1999) were recorded at multiple magnetic field strengths and temperatures. Recycle delays of 1.2 and 1.5 s were used at 500 and 750 MHz, respectively. Typically 12 to 14 spectra were recorded with refocusing fields, ν_CPMG_, between 50 and 850 Hz and two reference spectra (Mulder et al. 2001) were collected to convert measured peak intensities into relaxation rates. At 15 °C and 25 °C, experiments were collected at both 500 and 750 MHz. Experiments at 10 °C, 20 °C and 35 °C were recorded at 500 MHz only. Δloop-nDsbD, was studied at 25 °C at 750 MHz. The experiments were analyzed by global fitting using CATIA (Cpmg, Anti-trosy, and Trosy Intelligent Analysis, http://www.biochem.ucl.ac.uk/hansen/catia/) (Vallurupalli, Hansen, and Kay 2008, Baldwin, Hilton, et al. 2011, Alderson et al. 2019)using transition state theory to restrict the rates and linear temperature-dependent changes in chemical shifts when analyzing data across multiple temperatures (Vallurupalli, Hansen, and Kay 2008, Baldwin, Hilton, et al. 2011, Alderson et al. 2019).

The HSQC and HMQC pulse sequences developed by Skrynnikov et al. (Skrynnikov, Dahlquist, and Kay 2002) were used to determine the sign of the chemical shift differences between the major and minor states. Bruker versions of the sequences were kindly provided by Prof. L.E. Kay. Experiments were recorded in triplicate at 750 MHz using a spectrometer equipped with a cryoprobe. 48 scans were collected per ^15^N increment, using ^1^H and ^15^N sweep widths of 11904.762 and 2493.767 Hz, respectively, and with 1024 and 175 complex points in the ^1^H and ^15^N dimensions, respectively.

### Molecular dynamics simulations

MD simulations were run using GROMACS 4.5 (Van der Spoel et al. 2005). Simulations were started from the closed crystal structures of reduced nDsbD (nDsbD_red_) (PDB: 3PFU) (Mavridou et al. 2011) and oxidized nDsbD (nDsbD_ox_) (PDB 1L6P) (Goulding et al. 2002a). For each redox state of nDsbD, four simulations of 200 ns and ten simulations of 20 ns, all with different initial velocities, were run giving a combined duration of 1 μs for each protein. Trajectories were also initiated from open structures (PDB: 1VRS chain B) (Rozhkova et al. 2004); eight and ten 10 ns trajectories were run for nDsbD_red_ and nDsbD_ox_, respectively. Missing side-chain atoms were added using the WHAT-IF Server (https://swift.cmbi.umcn.nl/whatif) (Hekkelman et al. 2010). The histidine side chains were protonated and the cysteine side chains (C103 and C109) in the active site of nDsbD_red_ were represented as thiols; these choices were based on pH titrations monitored by NMR (Mavridou et al. 2012). The protein was embedded in rhombic dodecahedral boxes with a minimum distance to the box edges of 12 Å at NaCl concentrations of 0.1 M. Trajectories using a larger distance of 15 Å to the box edges showed no significant differences. The CHARMM 22 force field (MacKerell et al. 1998) with the CMAP correction (Mackerell, Feig, and Brooks 2004) and the CHARMM TIP3P water model was used. Electrostatic interactions were calculated with the Particle Mesh Ewald method (PME) (Essmann et al. 1995). The Lennard-Jones potential was switched to zero between 10 Å and 12 Å (Bjelkmar et al. 2010). The length of bonds involving hydrogen atoms was constrained using the PLINCs algorithm (Hess 2008). The equations of motion were integrated with a 2 fs time step. The simulation systems were relaxed by energy minimization and 4 ns of position-restrained MD in the NVT ensemble before starting the production simulations in the NPT ensemble at 25 °C and 1 bar for 10 ns, 20 ns or 200 ns. The Bussi-Donadio-Parinello thermostat (Bussi, Donadio, and Parrinello 2007), with a τ_T_ of 0.1 ps, and the Parinello-Rahman barostat (Parrinello and Rahman 1981), with a τ_P_ of 0.5 ps and a compressibility of 4.5 × 10^5^ bar^-1^, were used.

### Analysis of the MD simulations

The Visual Molecular Dynamics (VMD) program (Humphrey and Dalke 1996), GROMACS (Van der Spoel et al. 2005) and the MDAnalysis Python library (Michaud-Agrawal et al. 2011) were used to measure parameters including distances, torsion angles, accessibility and hydrogen bonds from the MD simulations. To validate our MD simulations, amide order parameters (S^2^) and residual dipolar couplings (RDCs) were calculated from them. Before calculating S^2^ and RDCs, we removed the overall tumbling from the simulations by aligning each frame to a reference structure. Order parameters were calculated in 5 ns blocks from the MD trajectories (Chandrasekhar et al. 1992) with S^2^ given by

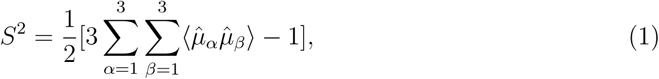

where α and β denote the x,y,z components of the bond vector μ.

To calculate RDCs, the principle axes of the reference structure were aligned with the experimentally determined alignment tensor. The use of a single reference structure is justified given the stability of the overall fold of nDsbD in the simulations. The RDC for a given residue in a given structure was then calculated using

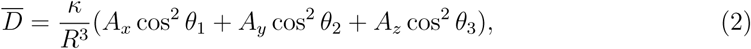

where κ depends on the gyromagnetic ratios γ_I_ γ_S_, the magnetic permittivity of vacuum μ_0_ and Planck’s constant and is defined as:

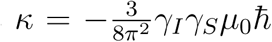

*R* is the bond length and is uniformly set to 1.04 Å. The angles θ_1_, θ_2_ and θ_3_ describe the orientation of the individual bonds with respect to the three principal axes of the alignment tensor. *A*_*x*_, *A*_*y*_ and *A*_*z*_ are the principal components of the alignment tensor. Errors in these S^2^ and RDC values were estimated using a bootstrap analysis using random sampling with replacement in 5 ns simulation blocks.

### Bioinformatics

2753 bacterial and archaeal representative proteomes from the NCBI database were searched for homologs of the N-terminal domain of *Escherichia coli* DsbD (nDsbD) (residues M1-V132) using blastp 2.2.28+ (NCBI) with the following cut-off values: evalue < 0.01, identity > 20%, coverage >50%. Hits were subsequently used as a blastp query against the *E. coli* MG1655 proteome (evalue < 0.01) and only the ones giving DsbD as best hit were retained. A full-length DsbD phylogenetic tree was built using fasttree 2.1.7 (Price, Dehal, and Arkin 2010) using the wag option. A subset of 731 sequences clustering around the *E. coli* DsbD were selected for subsequent analysis as a consistent phylogenetic group of proteins, mostly belonging to the β- and γ-proteobacteria. One sequence per bacterial genus was selected; this gave a set of 134 sequences which were aligned using muscle 3.8.31 (Edgar 2004). Percent conservation of individual positions was calculated using an in-house script.

### Crystallization, data collection, and structure determination of Δ loop-nDsbD_*ox*_

Crystals of Δloop-nDsbD_ox_ were obtained in vapour diffusion hanging drops containing a 1:1 mixture of Δloop-nDsbD_ox_ (10 mg/ml) in 20 mM Tris-HCl (pH 7.5), 50 mM NaCl and reservoir solution containing 28% w/v PEG 4000, 0.1 M ammonium sulphate and 0.1 M sodium acetate (pH 5.0); these conditions had previously yielded diffraction-quality crystals for nDsbD_red_ (Mavridou et al. 2011). The reservoir was sealed with vacuum grease and incubated at 16 °C. Crystals were cryoprotected by soaking for a few seconds in 20% v/v glycerol and 80% v/v reservoir solution, and frozen directly in the nitrogen stream of the X-ray apparatus. Datasets were collected with an oscillation angle of 162° (1°/frame) on a Rigaku RU-H3R rotating anode X-ray generator equipped with an R-AXIS IV image plate detector and an Oxford Cryosystems cryostream.

Diffraction data were processed and scaled with the Denzo/Scalepack package (Otwinowski and Minor 1997). Although datasets were initially collected to a resolution of 2.20 Å, completeness was low due to damage of the crystals because of ice formation. It was, therefore, decided to refine the data to 2.60 Å resolution to have satisfactory completeness (overall above 80%). The crystals belonged to space group P2_1_ (*a* = 37.33 Å, *b* = 81.34 Å, *c* = 46.39 Å and β = 100.66°). Five percent of reflections were flagged for R_free_ calculations. The structure was solved by molecular replacement with MOLREP (Murshudov, Vagin, and Dodson 1997) using the wild-type nDsbD_red_ structure (PDB accession code 3PFU) as a search model (Mavridou et al. 2011), and refined with Phenix.refine (Adams et al. 2010) and Coot (Emsley et al. 2010), which was used for manual fitting. The refinement converged at R = 25.2% and R_free_ = 29.2 %. Two protein molecules were found per asymmetric unit and 59 water molecules were also modelled. No electron density was observed before residue 10 and after residue 120, and before residue 9 and after residue 121 in chains A and B, respectively. The coordinates are deposited in the Protein Data Bank (PDB accession code 5NHI).

## Acknowledgements

L.S.S. acknowledges the Biotechnology and Biological Sciences Research Council (BBSRC) for a graduate studentship (BB/F01709X/1). D.G. acknowledges the Swiss National Science Foundation for the Postdoc Mobility (P300PA_167703) and Ambizione Fellowships (PZ00P3_180142). M.S.P.S. acknowledges funding from the Wellcome Trust (grant number 208361/Z/17/Z) and the BBSRC (grant number BB/R00126X/1). C.R. acknowledges funding from the Wellcome Trust (grant number 079440, and grant number 092532/Z/10/Z with S.J.F.). We thank Prof. D. Flemming Hansen for helpful discussions, Nick Soffe for help with the NMR pulse sequence programming, Prof. L.E. Kay for providing Bruker pulse programs for sign determination, and Albert Magnell for assistance with the expression and purification of Δloop-nDsbD. We are also grateful to all scientists involved in the review of this work for their constructive criticism. The authors would like to acknowledge the use of the University of Oxford Advanced Research Computing (ARC) facility in carrying out this work.

## Source Data and Figure Supplements

**Figure 2 – Source data 1. Quality factors (Q) and alignment tensor parameters (D**_**a**_, **R, θ, ϕ, Ψ) obtained from fits of experimental RDCs to X-ray structures for oxidized (1L6P) and reduced (3PFU) nDsbD and from analysis of the 1μs MD trajectories for oxidized and reduced nDsbD**.

| **nDsbD<sub>ox</sub> using the 1L6P X-ray structure** |  |  |  |  |  |  |  |
| --- | --- | --- | --- | --- | --- | --- | --- |
| **Region for which RDCs were fitted** | **Number of residues fitted** | **Q** | **$D_a$** | **R** | **$\theta$** | **$\phi$** | **$\psi$** |
| 8-122 | 87 | 0.34 | -15.7 | 0.56 | 83.6 | 86.5 | 70.2 |
| 12-122 | 83 | 0.26 | -16.4 | 0.61 | 83.1 | 85.1 | 71.0 |
| 12-122 w/o 106/108* | 81 | 0.24 | -16.8 | 0.61 | 83.7 | 84.8 | 71.7 |
| core | 46 | 0.21 | -17.7 | 0.61 | 84.7 | 83.2 | 72.9 |
| active site | 14 | 0.15 | -15.7 | 0.56 | 80.6 | 88.6 | 67.2 |
| **1 μs MD simulation of nDsbD<sub>ox</sub>** |  |  |  |  |  |  |  |
| 8-122 | 87 | 0.26 |  |  |  |  |  |
| **nDsbD<sub>red</sub> using the 3PFU X-ray structure** |  |  |  |  |  |  |  |
| 8-125 | 90 | 0.38 | -15.2 | 0.56 | 85.2 | 88.8 | 105.3 |
| 12-122 | 87 | 0.29 | -16.5 | 0.57 | 85.5 | 91.1 | 105.5 |
| 12-122 w/o 106/108* | 85 | 0.19 | -17.5 | 0.57 | 85.2 | 92.3 | 106.3 |
| core | 46 | 0.17 | -17.5 | 0.61 | 85.0 | 91.7 | 108.6 |
| active site | 16 | 0.16 | -18.6 | 0.52 | 88.4 | 93.9 | 106.2 |
| **1 μs MD simulation of nDsbD<sub>red</sub>** |  |  |  |  |  |  |  |
| 8-125 | 90 | 0.23 |  |  |  |  |  |

\*106/108 were excluded due to crystal contacts in the X-ray structures which may distort the conformation leading to poor fits; core means $\alpha$ 1, β-sandwich and tight turn(s); active site means $\alpha$ 2, β-strands but not loop (69-71).

**Figure 2 – Source data 2. Model-free analysis of** ^**15**^**N relaxation data for the cap loop and the flanking β-strands**.

|  | nDsbD<sub>ox</sub> |  |  | nDsbD<sub>red</sub> |  |  |
| --- | --- | --- | --- | --- | --- | --- |
| | $\{^1\text{H}\}\text{-}^{15}\text{N}$<br>NOE | $S^2$ | $\tau_e$ (ps) | $\{^1\text{H}\}\text{-}^{15}\text{N}$<br>NOE | $S^2$ | $\tau_e$ (ps) |
| V64 | 0.67 | 0.87 | 83 +/- 62 | 0.77 | 0.87 |  |
| W65 | 0.70 | 0.86 | 54 +/- 20 | 0.76 | 0.87 |  |
| H66 |  |  |  | 0.83 | 1.0 |  |
| E67 |  |  |  |  |  |  |
| D68 | 0.59 | 0.86 | 108 +/- 85 | 0.77 | 0.85 |  |
| E69 | 0.50 | 0.89 | 334 +/- 121 | 0.70 | 0.87 | 63 +/- 18 |
| F70 | 0.61 | 0.91 | 246 +/- 136 | 0.77 | 0.93 |  |
| Y71 | 0.63 | 0.88 | 99 +/- 61 | 0.75 | 0.93 |  |
| G72 | 0.69 | 0.83 | 47 +/- 14 | 0.74 | 0.85 |  |
| K73 |  |  |  | 0.72 |  |  |
| S74 | 0.76 | 0.93 |  | 0.80 | 0.94 |  |
| E75 | 0.72 | 0.84 |  | 0.72 | 0.87 |  |
| I76 | 0.84 | 0.81 |  |  |  |  |
| Y77 | 0.78 | 0.82 |  | 0.87 | 0.86 |  |

**Figure 6 – Source data 1. Parameters from a multi-temperature analysis of the relaxation dispersion data**. Data were acquired at five temperatures (10, 15, 20, 25, 35 °C) at 500 MHz and at two temperatures (15 and 25 °C) at 750 MHz. The data were analyzed globally (Figure 6 – figure supplement 1). Peaks for residues 63, 64, 65, 70, 71, 73 and 74 were sufficiently resolved for peak intensity data to be obtained in all experiments. At the extreme temperatures of 10 °C and 35 °C peak intensities could not be determined with confidence for residues 66, 68, 72 and 75 because of peak overlap or exchange broadening. Similarly, peak intensities for residue 69, at 10 °C and 15 °C could not be determined. In total 73 CPMG relaxation dispersion curves were entered into the global analysis. A minimum of two temperatures and two magnetic fields were available for each residue allowing robust parameters to be obtained. The data were analyzed as described previously (Baldwin, Hilton, et al. 2011) using CATIA (Vallurupalli, Hansen, and Kay 2008). In CATIA, magnetization is simulated to match the experiment as closely as possible. To this end, the peak positions are entered into the program as well as the carrier frequency, allowing off resonance effects of the 180° pulses to be accounted for. The difference between in-phase and anti-phase relaxation rates of coherences formed during the experiment was assumed to be 0.2 s^-1^ (Vallurupalli, Hansen, and Kay 2008). The populations of each state and rates of interconversion were assumed to follow Arrhenius behavior and chemical shifts were assumed to have a linear temperature dependence.

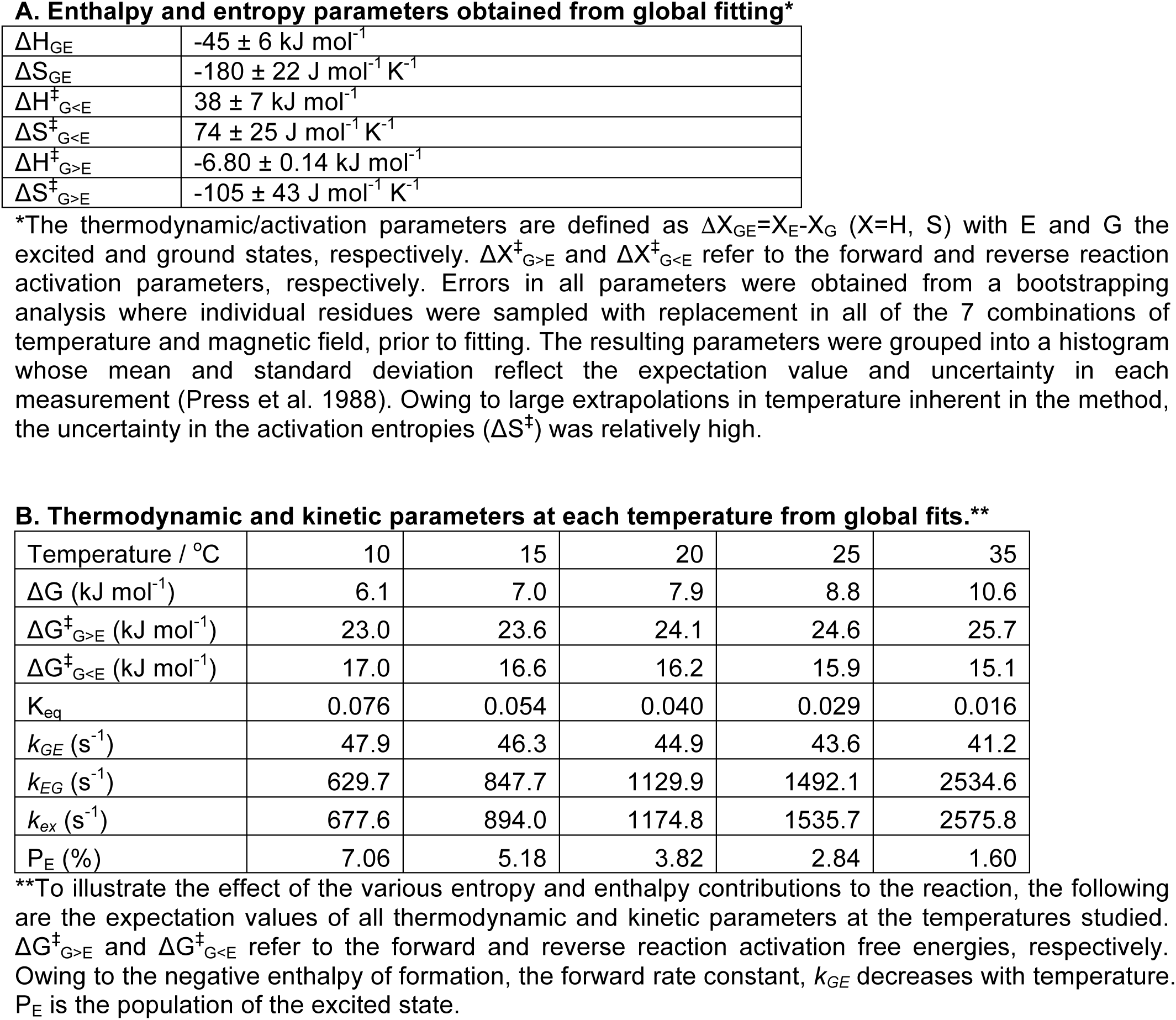
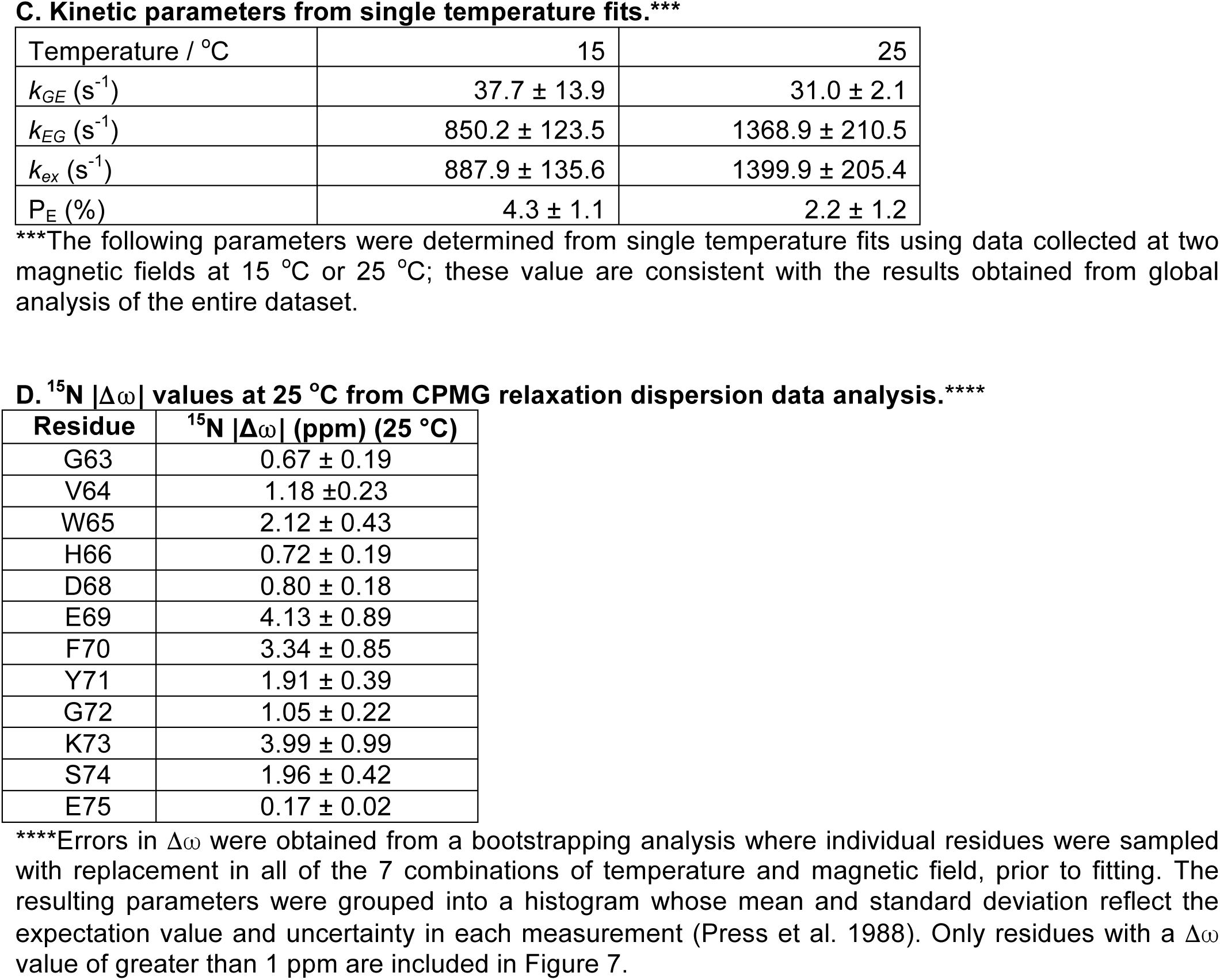

**Figure 6 – Source data 2. Crystallographic data collection and refinement statistics for Δloop-nDsbD**_**ox**_

| **Unit Cell** |  |
| --- | --- |
| Space Group | P2<sub>1</sub> |
| Cell dimensions (Å) | $a = 37.33, b = 81.34, c = 46.39, \beta = 100.66^\circ$ |
| **Data Collection** |  |
| Temperature (K) | 100 |
| Wavelength (Å) | 1.5418 |
| Resolution (Å) | 39.77 to 2.60 (2.74 to 2.60)* |
| $\langle I/\sigma \rangle$ | 5.4 (3.3)* |
| R<sub>meas</sub> | 0.31 (0.38)* |
| Multiplicity | 3.1 (2.9)* |
| **Refinement** |  |
| Refinement program | Phenix.refine |
| Number of unique reflections used | 6808 |
| Completeness (%) | 80.7 (79.0)* |
| R-factor (%) | 25.2 |
| R<sub>free</sub> (%) | 29.2 |
| Average B (Å<sup>2</sup>) | 26.0 |
| r.m.s.d. bond lengths (Å) | 0.003 |
| r.m.s.d. bond angles (°) | 0.721 |
| **Ramachandran** |  |
| Favoured | 211 (95.0%) |
| Allowed | 9 (4.1%) |
| Outliers | 2 (0.9%) |
| **Asymmetric unit contents** |  |
| Polypeptide chains | 2 |
| No. of atoms: total/non-solvent | 1838/1779 (13 in alternative conformations) |

\* values in parentheses are for corresponding highest resolution cell
r.m.s.d. = root mean square deviation

**Figure 7 – Source data 1. Determination of the sign of the** ^**15**^**N chemical shift difference between the excited state and the ground state determined by comparison of HSQC and HMQC spectra collected at 750 MHz**.**\***

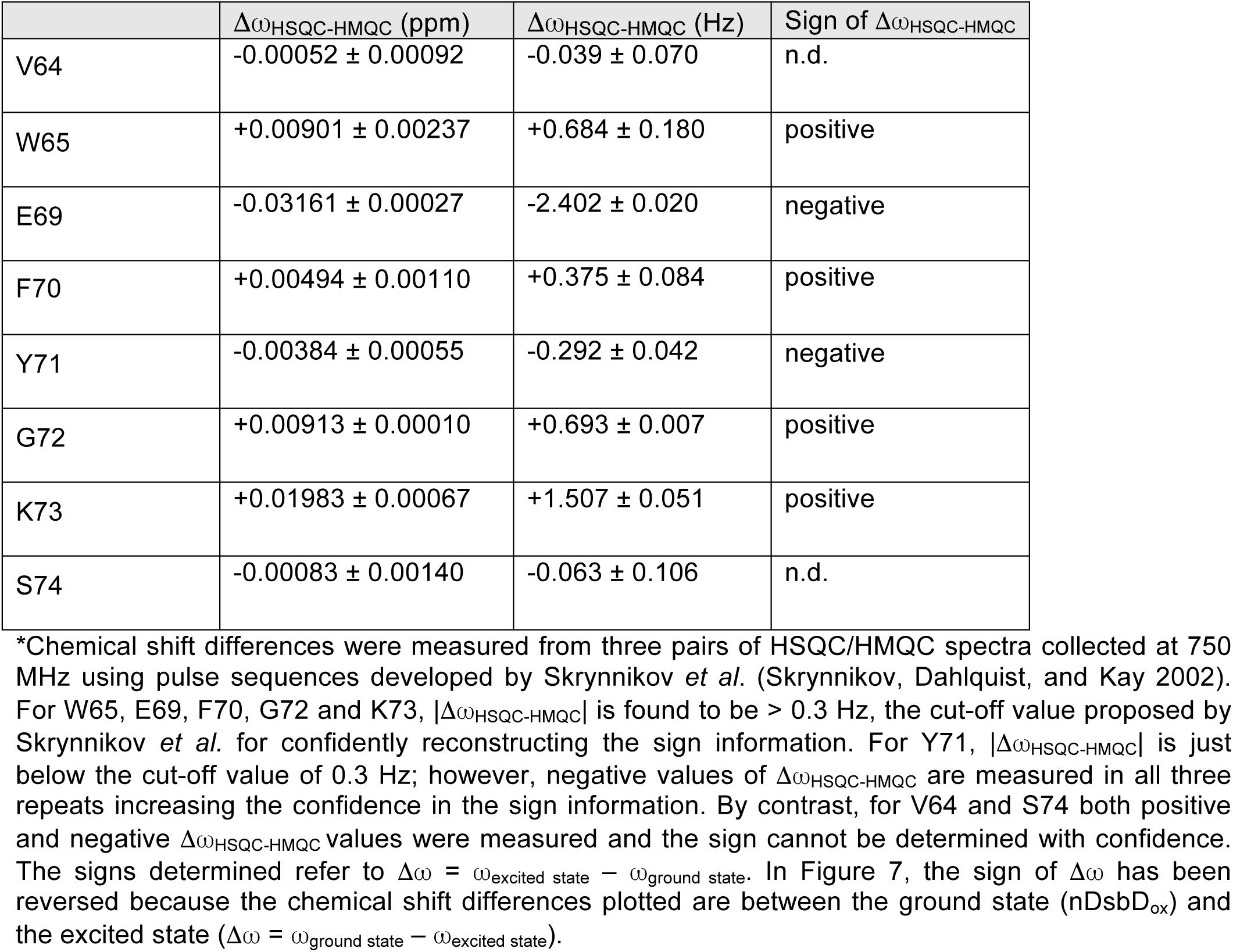

**Figure 2 – figure supplement 1.**
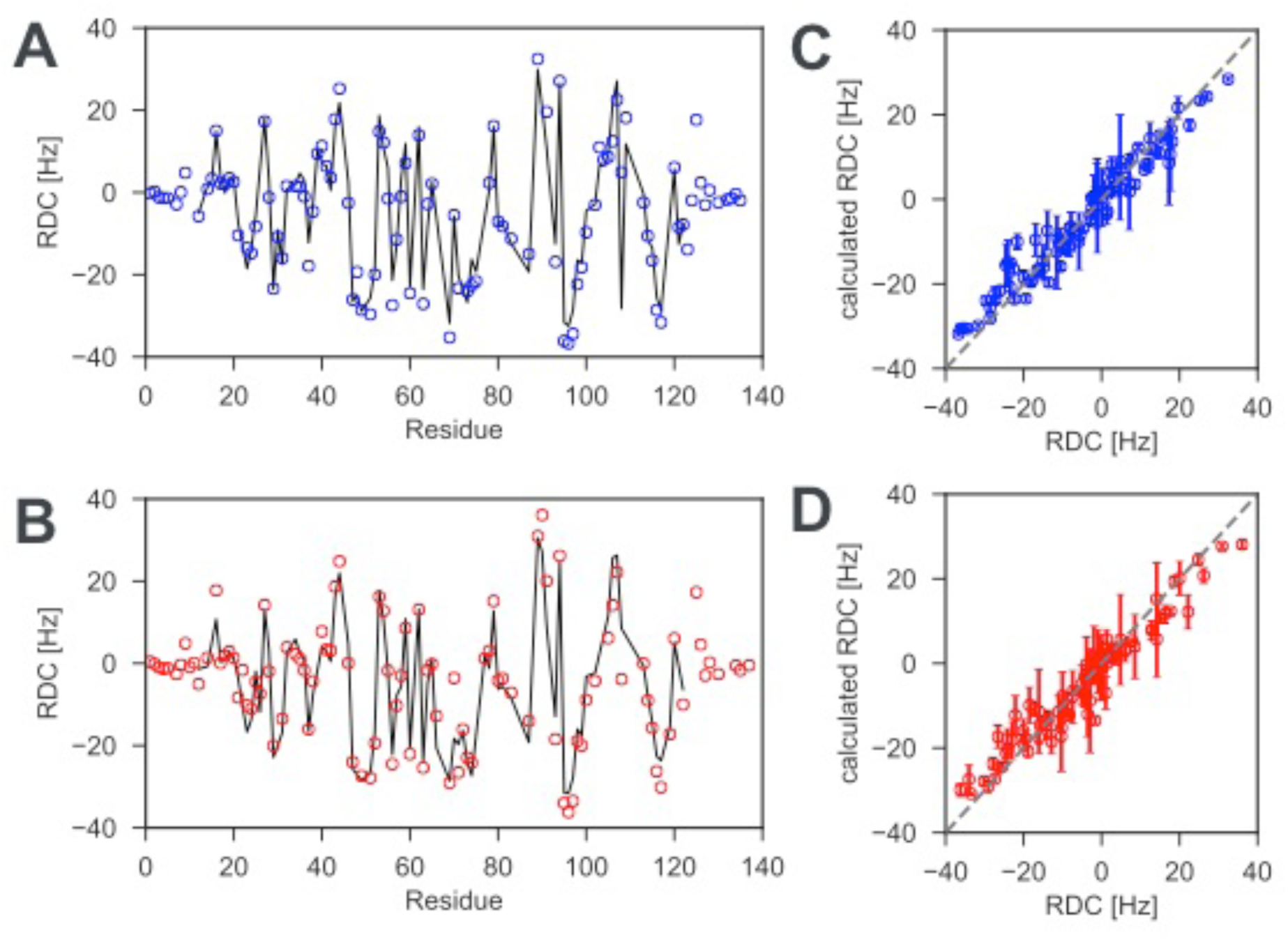
^1^H -^15^N residual dipolar couplings (RDCs) for reduced and oxidized nDsbD. To probe the solution structure of nDsbD, ^1^H-^15^N RDCs, which are sensitive reporters of protein structure, were measured for (A) reduced and (B) oxidized nDsbD. Experimental RDCs are indicated by the open circles while RDCs calculated by fitting the alignment tensor to the X-ray structure (PDB:3PFU for reduced and PDB:1L6P for oxidized nDsbD) are indicated by the solid line. For most residues very similar RDCs were measured for reduced and oxidized nDsbD. The small RDCs measured for residues 1-8 and 126-135 in both redox states are likely to indicate conformational dynamics that lead to averaging of these RDCs; values for most of these residues cannot be predicted from the X-ray structures because electron density is not observed before residue 8 and after residue 125 in the structures. By contrast, the RDCs for the core β-sandwich are accurately predicted by the X-ray structures; Q values of 0.17 and 0.21 are obtained for reduced and oxidized nDsbD, respectively (Figure 2 – source data 1). When fits are carried out for residues 12-122, which includes the core β-sandwich and the active-site/cap-loop residues, Q values of 0.19 and 0.24 are obtained for reduced and oxidized nDsbD, respectively (Figure 2 – source data 1). The similar Q values for the two sets of RDCs suggest that the orientation of the active-site/cap-loop residues relative to the core β-sandwich of both reduced and oxidized nDsbD in solution is described well by the X-ray structures in which the cap loop adopts a closed conformation. Experimental RDCs are compared with RDCs calculated from MD simulations for (C) reduced and (D) oxidized nDsbD. For both nDsbD_red_ and nDsbD_ox_, the agreement between calculated and experimental RDCs improved for the 1μs MD ensemble compared to the static X-ray structures. It is also interesting to note that the RDCs for residues 106 and 108, which gave poor agreement in the fits to the static X-ray structures due to crystal contacts, agree well with the values predicted from the MD simulations. Errors in RDCs calculated from the MD simulations were determined using a bootstrap analysis.

**Figure 2 – figure supplement 2.**
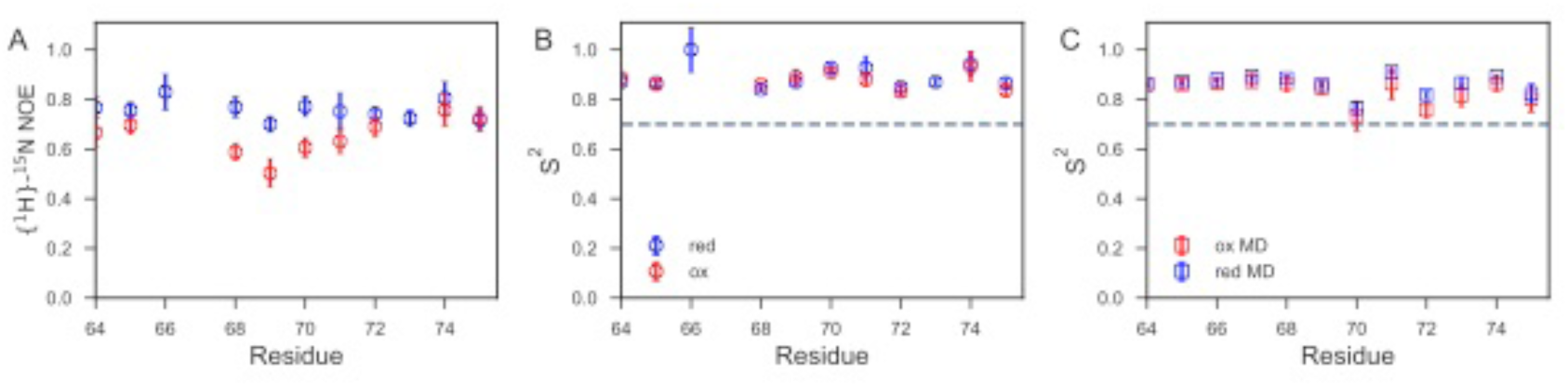
Fast time-scale dynamics of the cap loop. (A) Comparison of experimental {^1^H}-^15^N heteronuclear NOE values for nDsbD_red_ (blue) and nDsbD_ox_ (red). (B) Comparison of order parameters (S^2^) derived from model-free analysis of ^15^N relaxation data for nDsbD_red_ (blue) and nDsbD_ox_ (red). (C) Comparison of order parameters (S^2^) derived from the MD simulations for nDsbD_red_ (blue) and nDsbD_ox_ (red). Errors in the experimental {^1^H}-^15^N heteronuclear NOE and the S^2^ order parameters were determined as described previously using Monte Carlo error estimation (Smith et al. 2013).

**Figure 2 – figure supplement 3.**
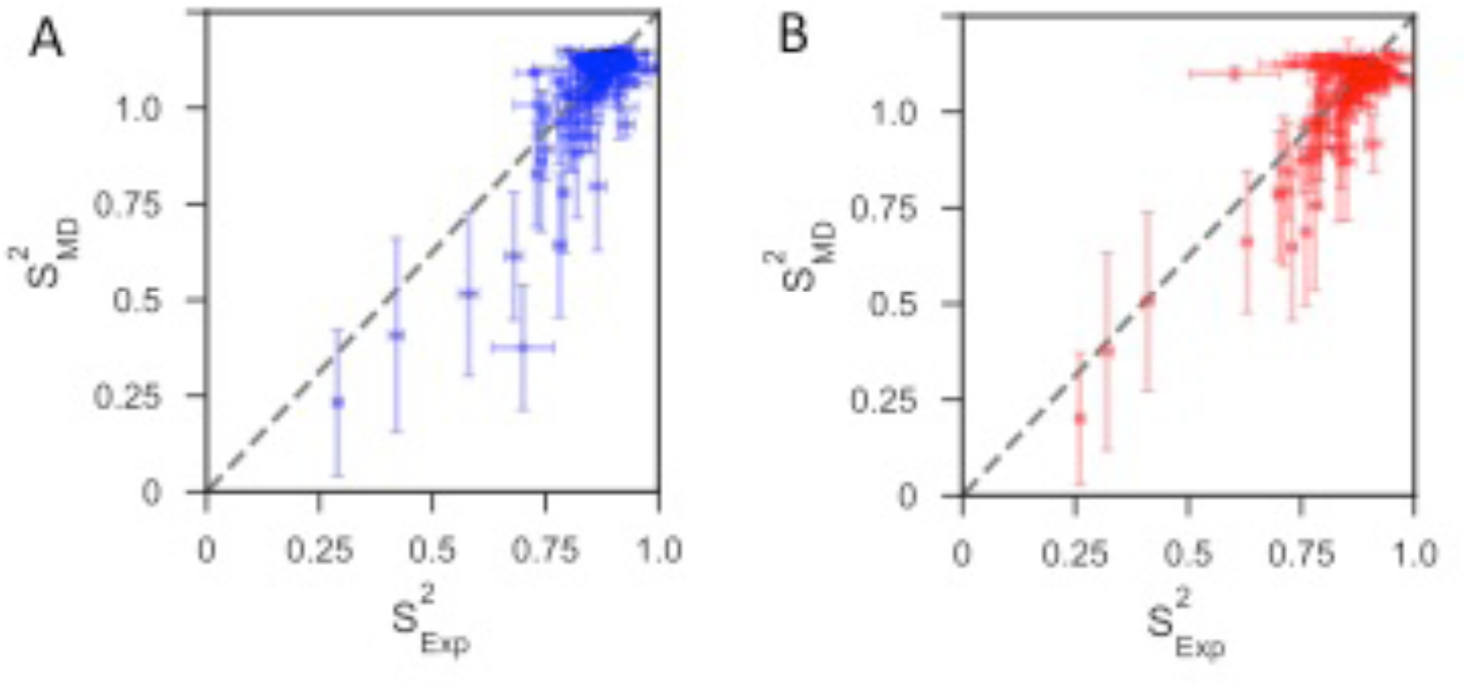
Comparison of backbone order parameters (*S*^*2*^) derived from experimental ^15^N relaxation data and calculated from the 1μs MD simulations for (A) reduced and (B) oxidized nDsbD. The extent of the fast-time scale (ps-ns) dynamics observed experimentally by NMR is on the whole correctly reproduced by the MD simulations. The root mean-square deviation (RMSD) between the order parameters detemined from model-free analysis and from the simulations is 0.08 for both reduced and oxidized nDsbD. The N- and C-termini were very flexible, but most residues in the folded segments of nDsbD were well ordered with S^2^ > 0.80, which agrees with experiment. Errors in S^2^ calculated from the MD simulations were determined using a bootstrap analysis.

**Figure 3 – figure supplement 1.**
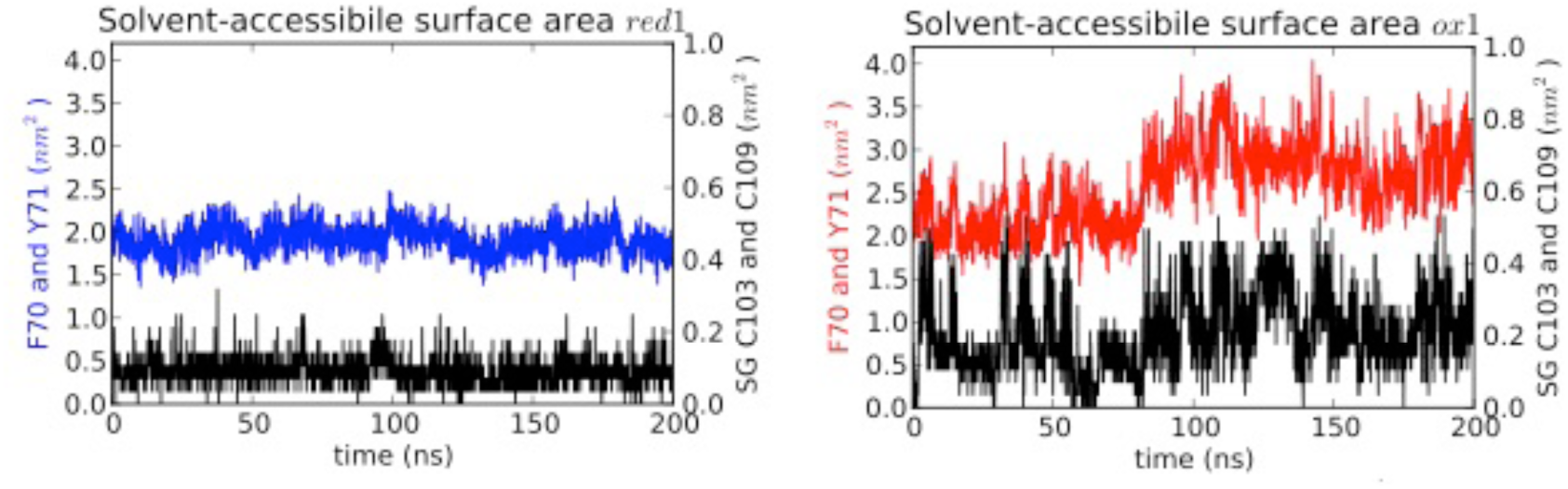
Solvent accessible surface area (SASA) measured for MD simulations. SASA is plotted for the aromatic cap loop residues F70 and Y71 and for the Sγ of C103 and C109 in simulations of nDsbD_red_ (left) and nDsbD_ox_ (right). The Sγ of C103 and C109 remain largely shielded from solvent in nDsbD_red_ but become more exposed during loop opening in nDsbD_ox_. The SASA of the aromatic rings of F70 and Y71 also increases during the main loop opening event between 80 and 200 ns. These data are derived from the trajectories shown in Figure 3A and 3F.

**Figure 3 – figure supplement 2.**
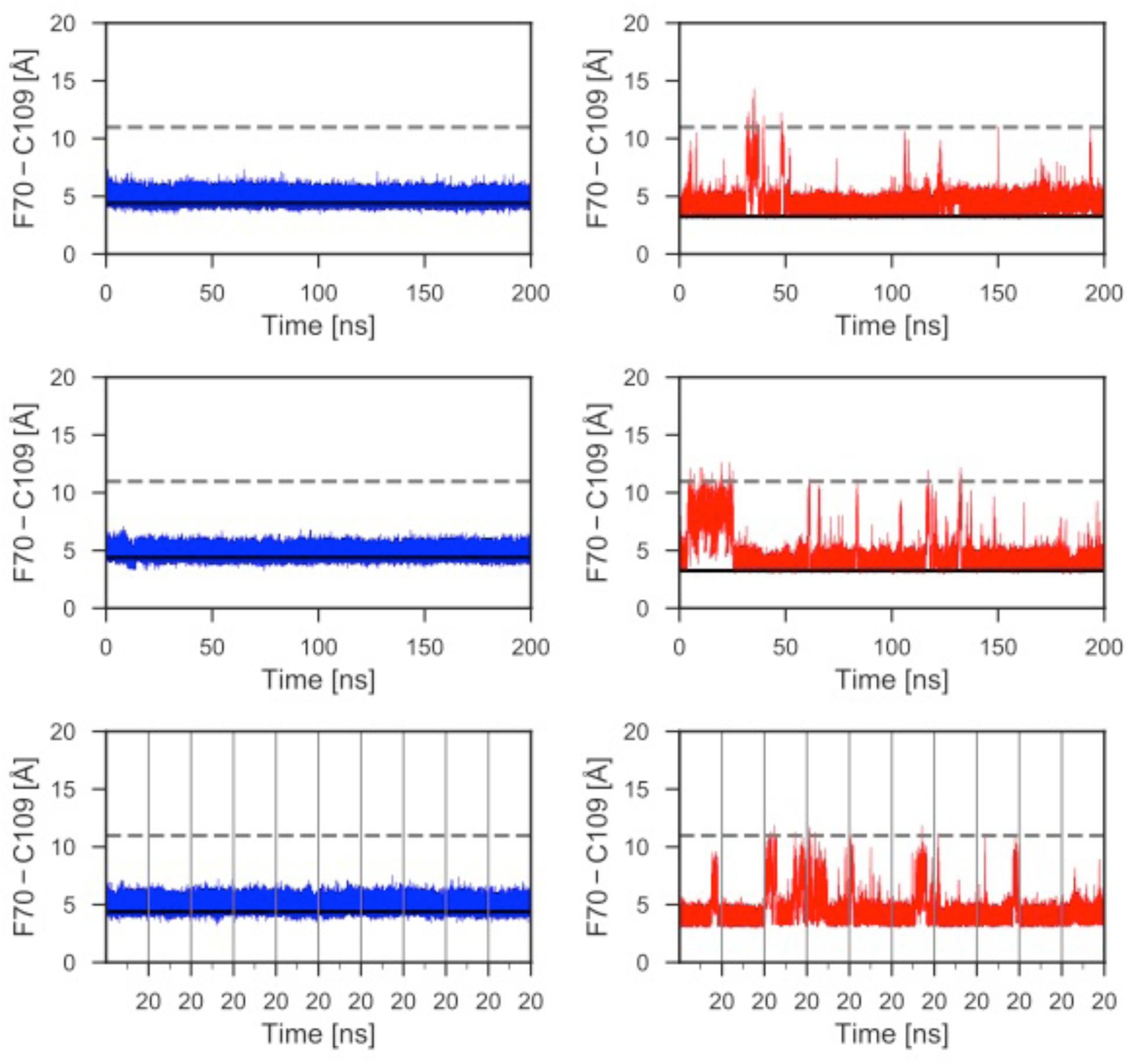
Distances between the aromatic ring of F70 and the Sγ of C109 for additional MD simulations. Data for an additional 600 ns of MD simulations, not illustrated in Figure 3, are shown for reduced (left) and oxidized (right) nDsbD. The conformational state of the cap loop was followed by tracking the distance between the centre of the aromatic ring of F70 and the sulfur atom of C109. The solid and dashed lines at ∼3-4 and ∼11 Å show the F70 ring to C109 Sγ distance in the closed (3PFU/1L6P) and open (1VRS) X-ray structures, respectively. The cap loop remains stably closed in simulations of nDsbD_red_ (left). The cap loop conformation is more flexible in simulations of nDsbD_ox_ (right) showing several transient and longer-lived opening events. Data for ten 20 ns simulations are combined in the bottom panels; the vertical grey lines show the starting and ending points of these shorter 20 ns simulations.

**Figure 4 – figure supplement 1.**
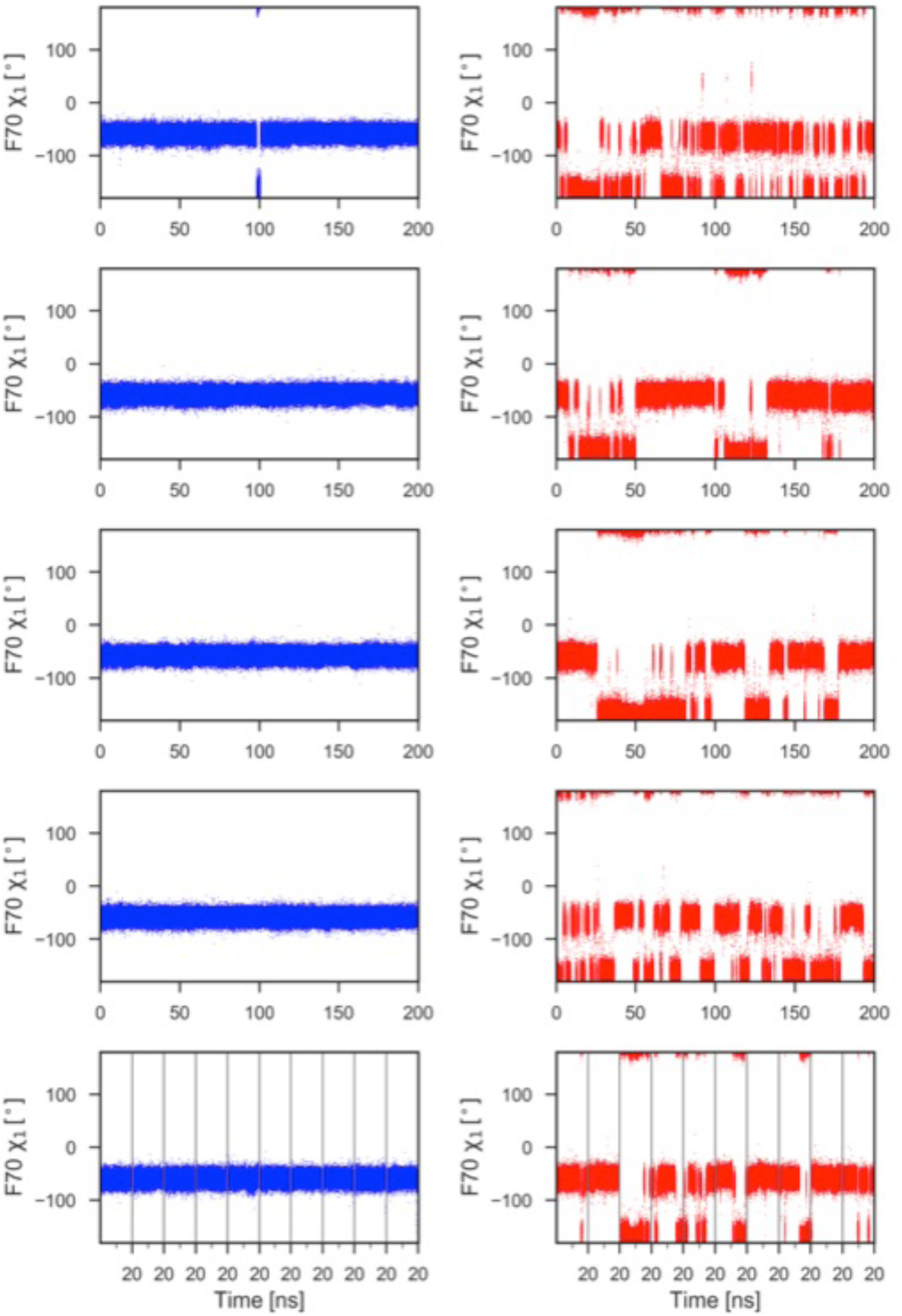
F70 χ _1_ torsion angle during 1μs of MD simulations for reduced (left) and oxidized (right) nDsbD. With the exception of a brief excursion to the trans (t) conformer (χ_1_ = -180°) in the top left panel, F70 remains in the gauche- (g-) conformer (χ1 = -60°) throughout the 1μs of MD simulations in nDsbD_red_ (left panels). By contrast, in nDsbD_ox_, F70 interconverts frequently between the g- and t conformers (right panels). In closed structures (F70-C109 distance < 6 Å), the g- and t conformers are sampled 61.5% and 38.5% of the time, respectively, while in open structures the g- and t conformers are sampled 52.9% and 47.1% of the time, respectively. Data for ten 20 ns simulations are combined in the bottom panels; the vertical grey lines show the starting and ending points of these shorter 20 ns simulations.

**Figure 4 – figure supplement 2.**
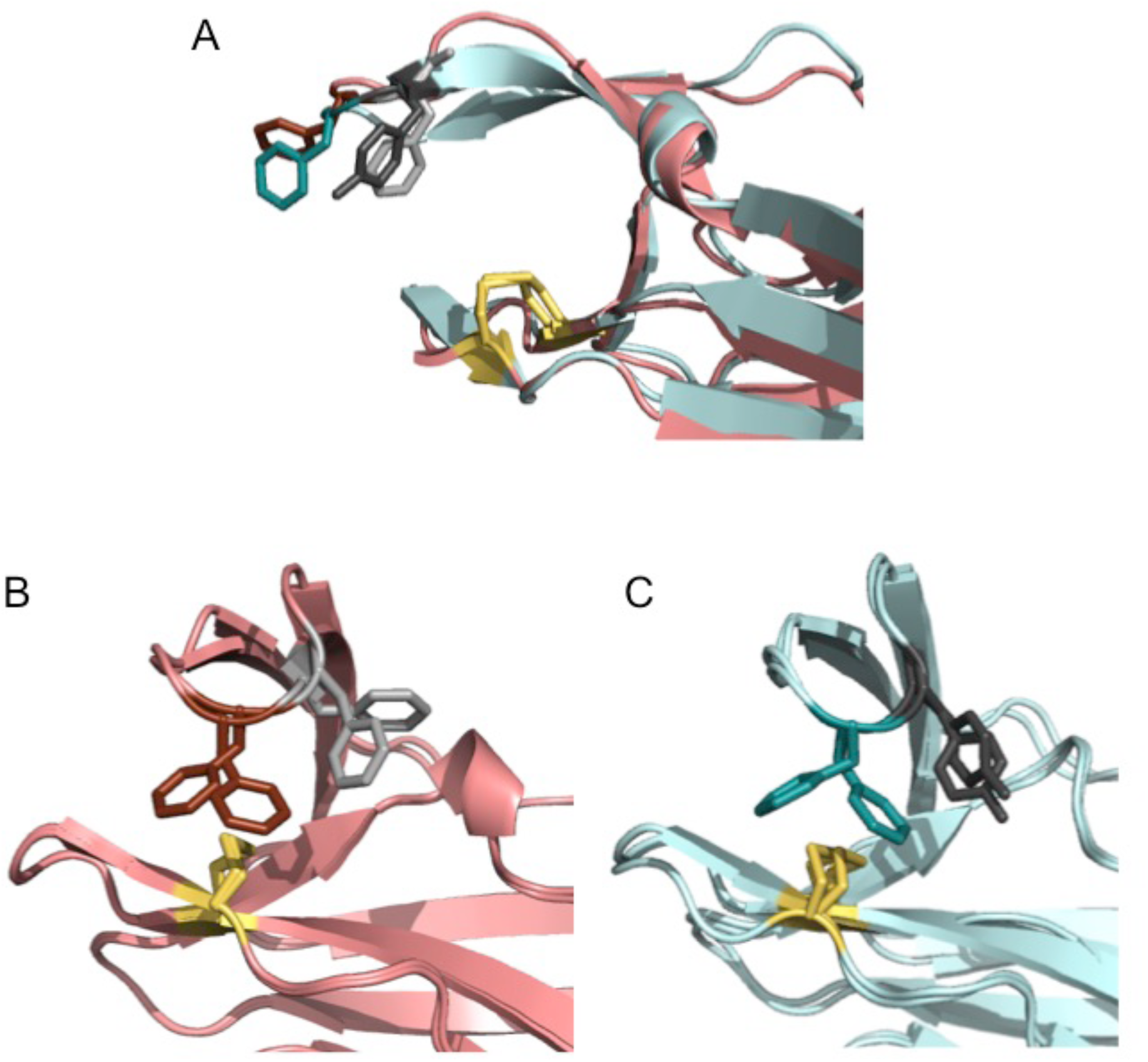
Comparison of ‘snapshots’ of *E. coli* nDsbD_ox_ from MD simulations with the X-ray structure of *Neisseria meningitidis* nDsbD_ox_. *N. meningitidis* nDsbD_ox_ crystallizes in the P2_1_3 space group with 6 molecules (denoted *A* to *F*) in the asymmetric unit (PDB: 6DPS (Smith et al. 2018)). (A) One of the six molecules (*E*) in the asymmetric unit has an open “Phen cap” loop. This structure superimposes well with the open conformation of *E. coli* nDsbD_ox_ observed in MD simulations. (B) The other 5 molecules in the *N. meningitidis* nDsbD_ox_ asymmetric unit have a closed “Phen cap” loop with the aromatic ring of F66 packing onto the C100-C106 disulfide bond in both gauche- and trans conformations. Shown here is a superposition of molecules *B* and *C* from the unit cell. (C) Superposition of two closed conformations of *E. coli* nDsbD_ox_ observed in the MD simulations showing F70 adopting the ‘frustrated’ gauche- and trans orientations. (A-C) The backbone of nDsbD_ox_ from *E. coli* and *N. meningitidis* are shown in cyan and salmon, respectively. F70 (dark teal), Y71 (dark grey), F66 (dark salmon), F67 (light grey) and the C103-C109 and C100-C106 disulfide bonds (yellow) are shown in a stick representation.

**Figure 5 – figure supplement 1.**
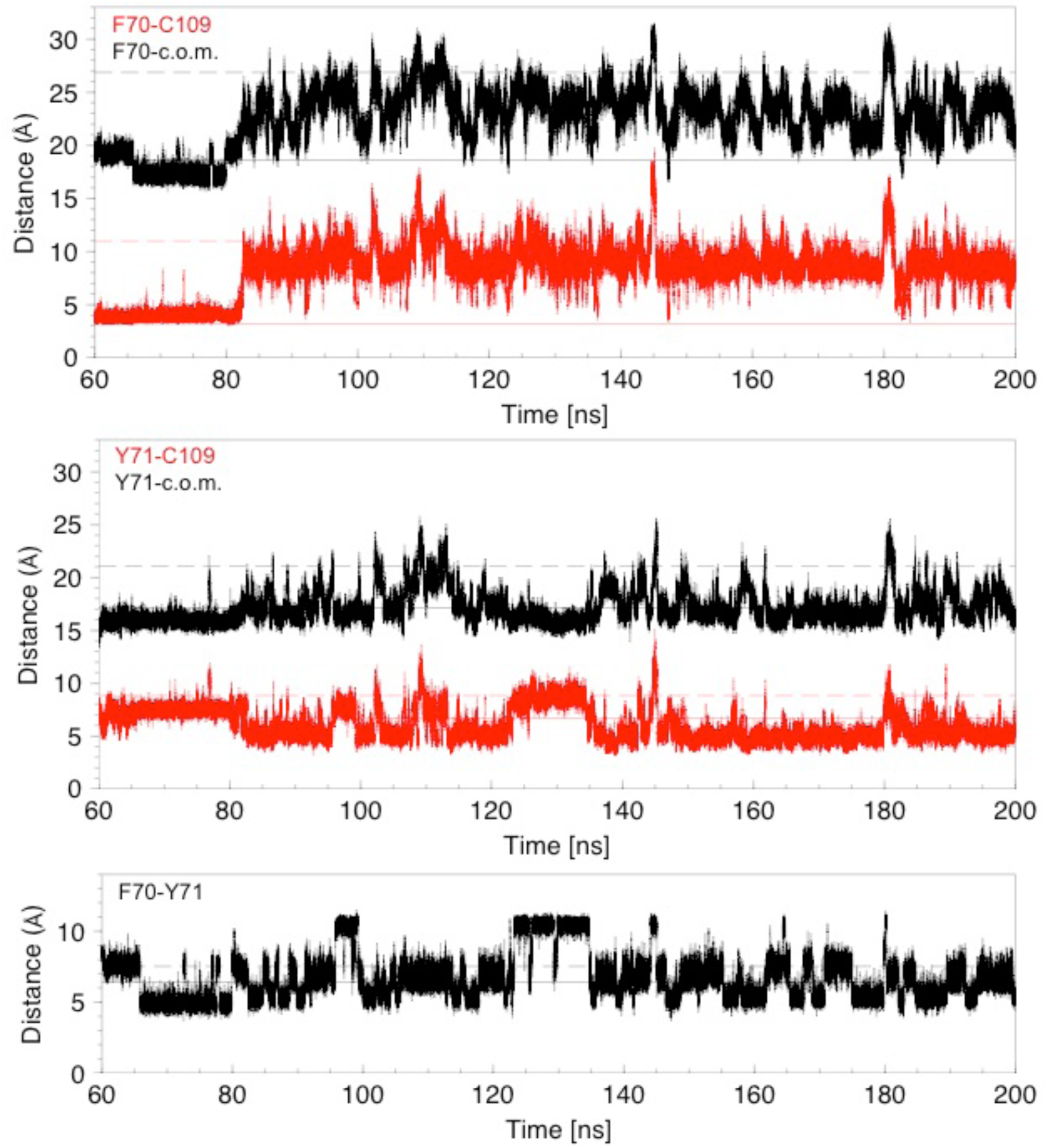
Analysis of distances for 140 ns of a molecular dynamics trajectory showing cap loop opening. Distances are derived from the MD trajectory shown in Figure 3F. Top panel: The distance between the centre of the F70 aromatic ring and C109 Sγ (red) or the centre of mass (c.o.m.) (black) are plotted as a function of time. The solid and dashed lines in the three panels show these distances in the closed 1L6P and open 1VRS X-ray structures. The centre of mass of nDsbD in the 1L6P X-ray structure was determined to coincide with W33 Cε2. Both these distances are a good indicator of whether the cap loop is closed or open. The distance to the centre of mass decreases at ∼66 ns because of the change of the F70 χ1 conformation from gauche- to trans. Middle panel: The distance between the centre of the Y71 aromatic ring and C109 Sγ (red) or the centre of mass (c.o.m) (black) are plotted as a function of time. The distance from Y71 to C109 is a less clear indicator of cap loop opening because this distance first decreases as Y71 moves past C109 and then increases again as Y71 adopts more open conformations. In structures in which Y71 sits between F70 and C109, and blocks loop closing, the distance of Y71 to C109 is reduced. Bottom panel: The distance between the centres of the aromatic rings of F70 and Y71 is plotted as a function of time. This can vary from ∼5 to ∼11 Å. The shorter distance usually corresponds to F70 χ1 in the trans conformation. The longer distance, observed at ∼95-100 and ∼125-135 ns, corresponds to the rings pointing in opposite directions usually with the F70 and Y71 χ1 in the gauche- and trans conformations, respectively.

**Figure 5 – figure supplement 2.**
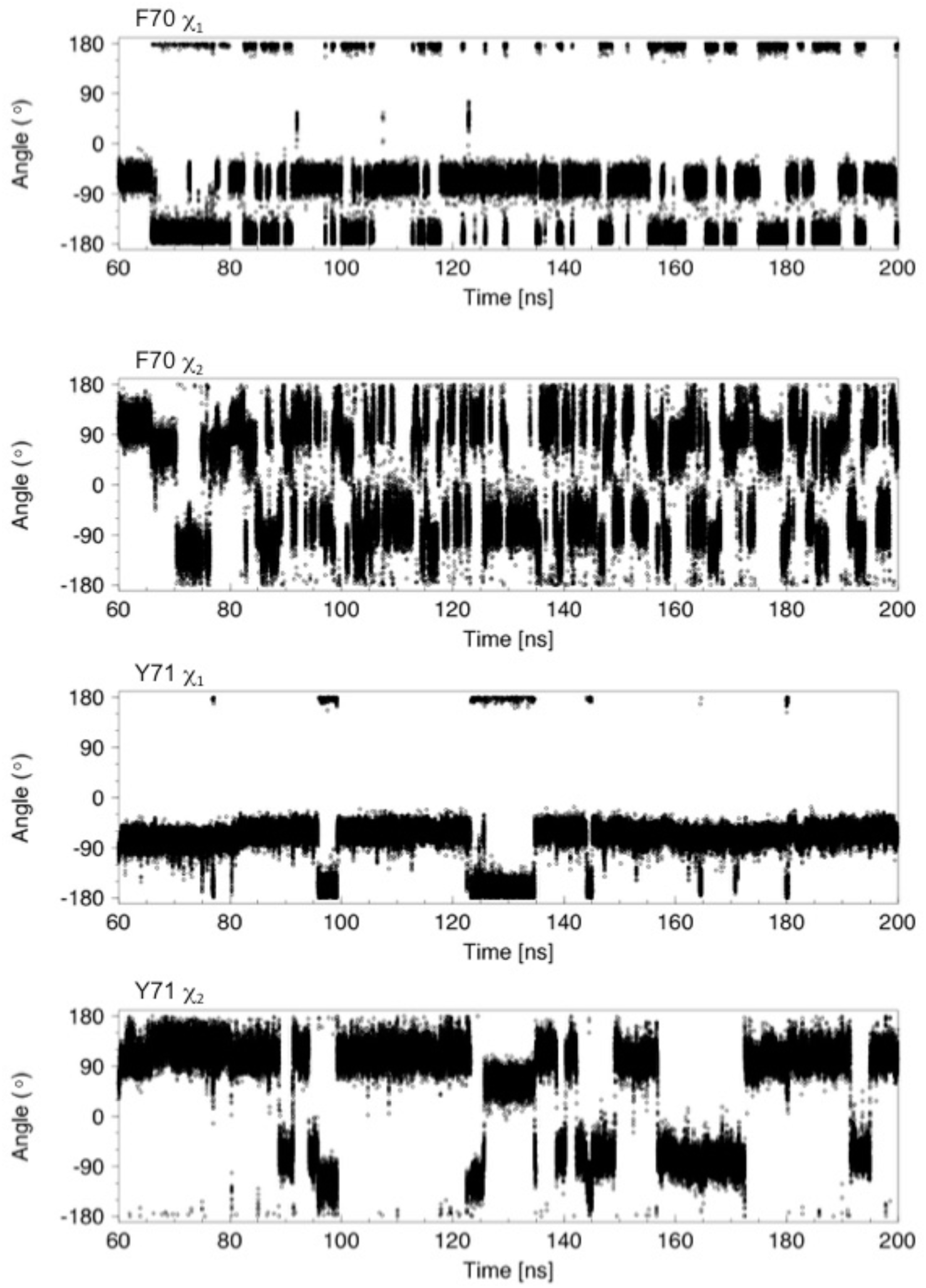
Analysis of χ 1 and χ 2 torsion angles for 140 ns of a molecular dynamics trajectory showing cap loop opening. The sidechain χ1 and χ2 torsion angles of F70 and Y71 are plotted as a function of time. In nDsbD_ox_, F70 interconverts between the gauche- (−60°) and trans (−180°) conformations. At 60 ns, the cap loop is closed with F70 χ1 in the gauche- conformation. At ∼66 ns, F70 χ1 switches to the trans conformation; this is accompanied by a change in the F70 χ2 angle from ∼120° to ∼65°. The cap loop begins to open at 82.5 ns and this coincides with a change of the F70 χ1 from -60° to -180°. After loop opening, F70 χ1 continues to interconvert between the gauche- and trans conformations. In the more exposed open conformation the aromatic ring of F70 undergoes frequent ring flips, as evidenced by 180° changes in χ2, but also changes between the ∼120° (or ∼ -60°) to ∼65° (or ∼ -115°) orientations observed before loop opening. The Y71 χ1 torsion angle changes less frequently than F70 and has a marked preference for the gauche-conformer. Transitions to the trans conformer correlate with the largest F70-Y71 ring-to-ring distances observed in panel (A); these changes to the trans conformer are coupled to changes in χ2 from ∼120° (or ∼ -60°) to ∼65° (or ∼ -115°). Ring flips of Y71 are less frequent due to its less exposed conformation.

**Figure 5 – figure supplement 3.**
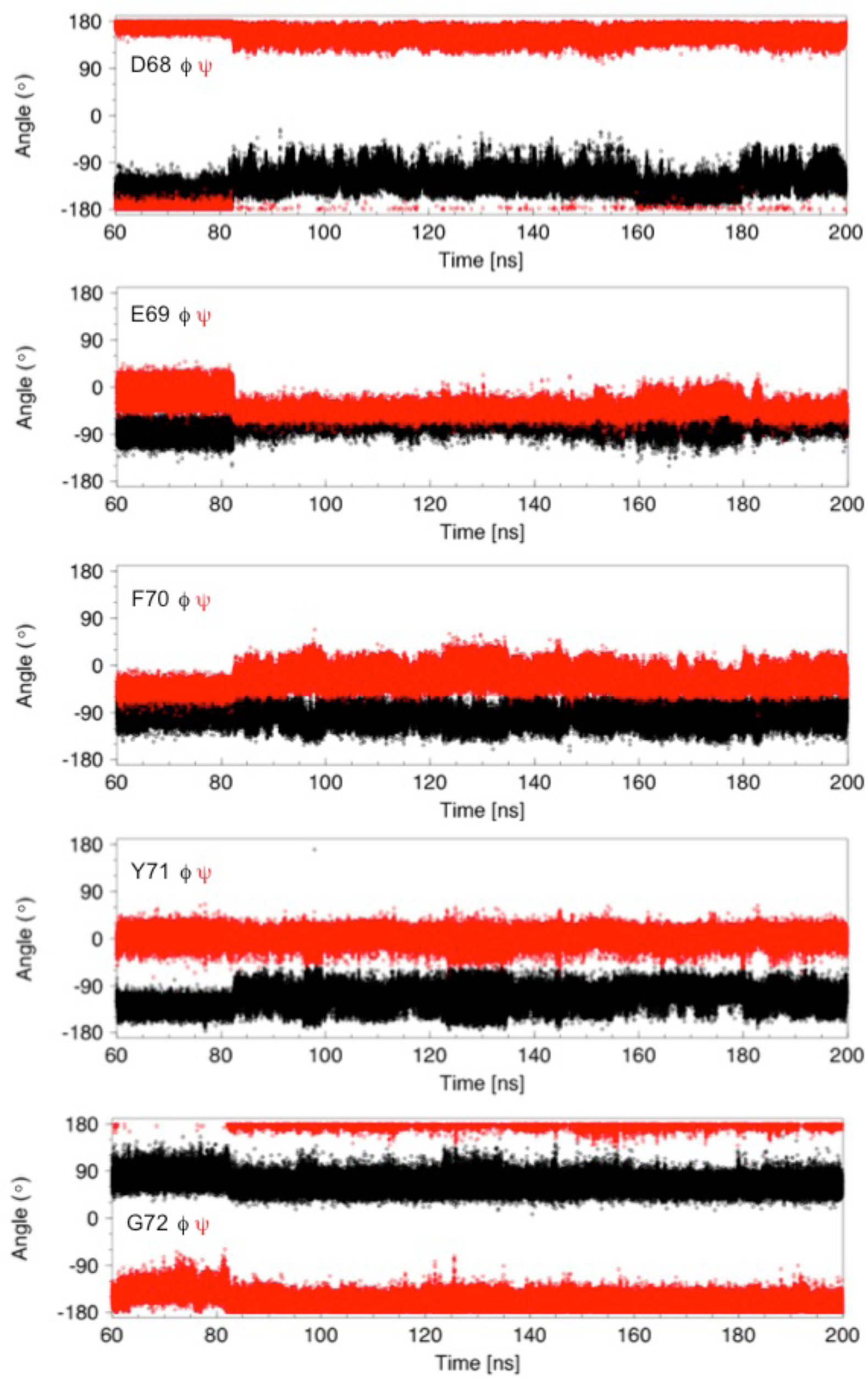
Analysis of ϕ and Ψ torsion angles for 140 ns of a molecular dynamics trajectory showing cap loop opening. Backbone ϕ (black) and Ψ (red) torsion angles are plotted as a function of time for residues D68, E69, F70, Y71 and G72. These angles change at 82.5 ns, the time at which the cap loop opens. The ϕ and Ψ values for E69 become less variable in the open loop while the values for the other residues appear to be more variable. Between ∼160 and 180 ns, the ϕ and Ψ angles of D68 and E69 return to values observed when the loop is closed. These changes are correlated with the presence (in the closed structure) or absence (in most of the open conformations) of a hydrogen bond between the backbone HN of Y71 and the D68 sidechain carboxyl group.

**Figure 5 – figure supplement 4.**
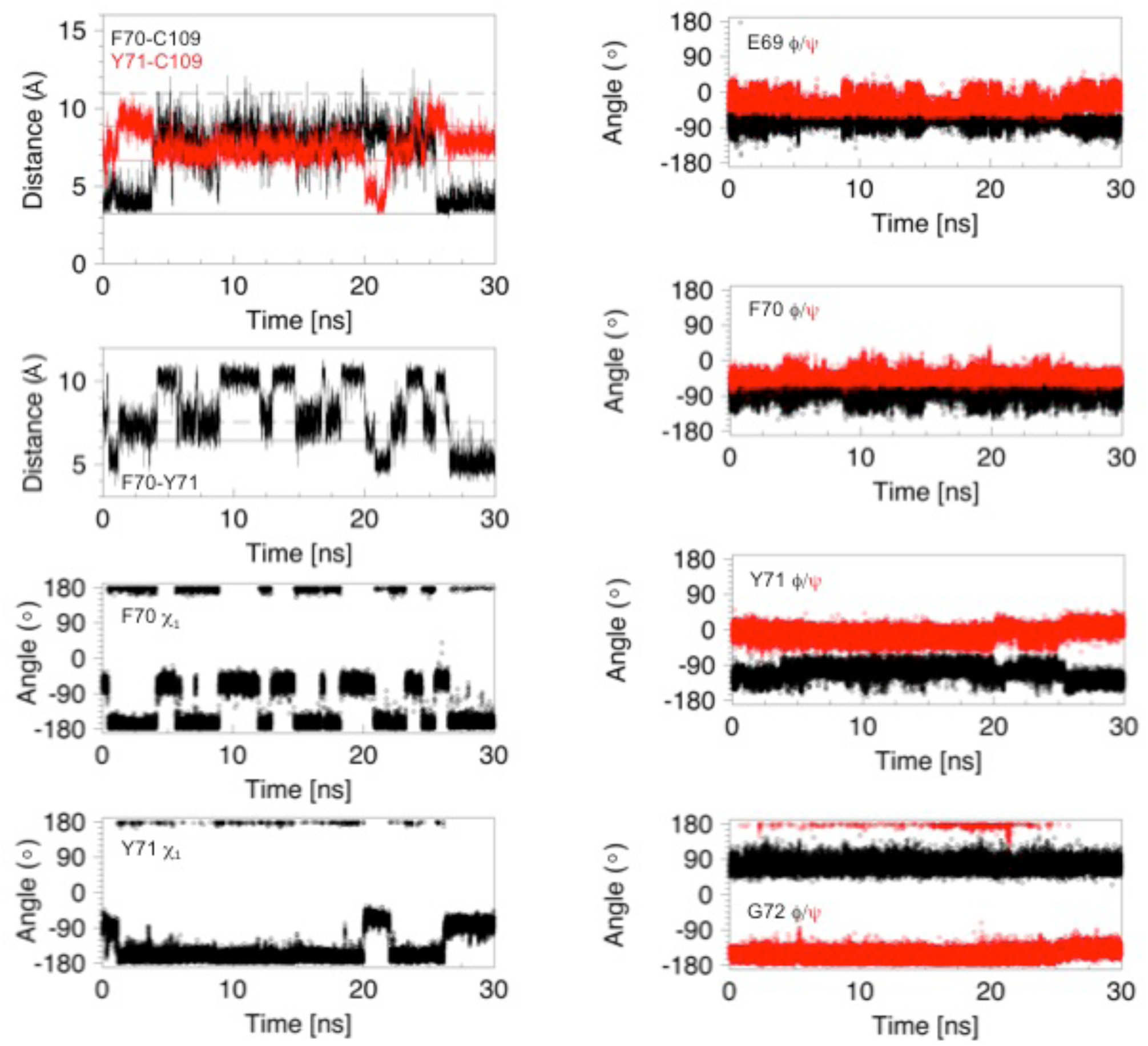
Analysis of distances and torsion angles for 30 ns of a molecular dynamics trajectory showing a 22 ns cap loop opening. Structural parameters, derived from the first 30 ns of the MD trajectory shown in Figure 3 – figure supplement 2 (middle right panel), are plotted as a function of time. The four panels on the left show the distance between C109 Sγ and the centres of the F70 (black) and Y71 (red) aromatic rings, the distance between the centre of the aromatic rings of F70 and Y71, and the χ1 torsion angles of F70 and Y71. The four panels on the right show the ϕ (black) and Ψ (red) torsion angles for E69, F70, Y71 and G72. The cap loop opens after 3.8 ns and remains open for 21.7 ns. The loop opens with F70 in the trans conformer but this χ1 angle changes several times while the loop is open. Before loop opening, the aromatic ring of Y71 is farther away from C109 than in the 1L6P X-ray structure. This is due to the trans χ1 conformer. At ∼20 ns the Y71 χ1 switches to the gauche-conformer; this brings the ring of Y71 closer to C109 and places the ring in between F70 and C109 in a ‘blocking’ conformation. At ∼22 ns the ring of Y71 moves out of this position, and away from C109, allowing the loop to close at 25.5 ns. The backbone torsion angles of E69, F70, Y71 and G72 show some of the features in the open conformations that have been highlighted for the major opening event in Figure 5 – figure supplement 3. When the loop closes at 25.5 ns, these torsion angles return to values typical of the closed conformation of nDsbD. Snapshots of structures from this opening event are shown in Figure 5 – figure supplement 5.

**Figure 5 – figure supplement 5.**
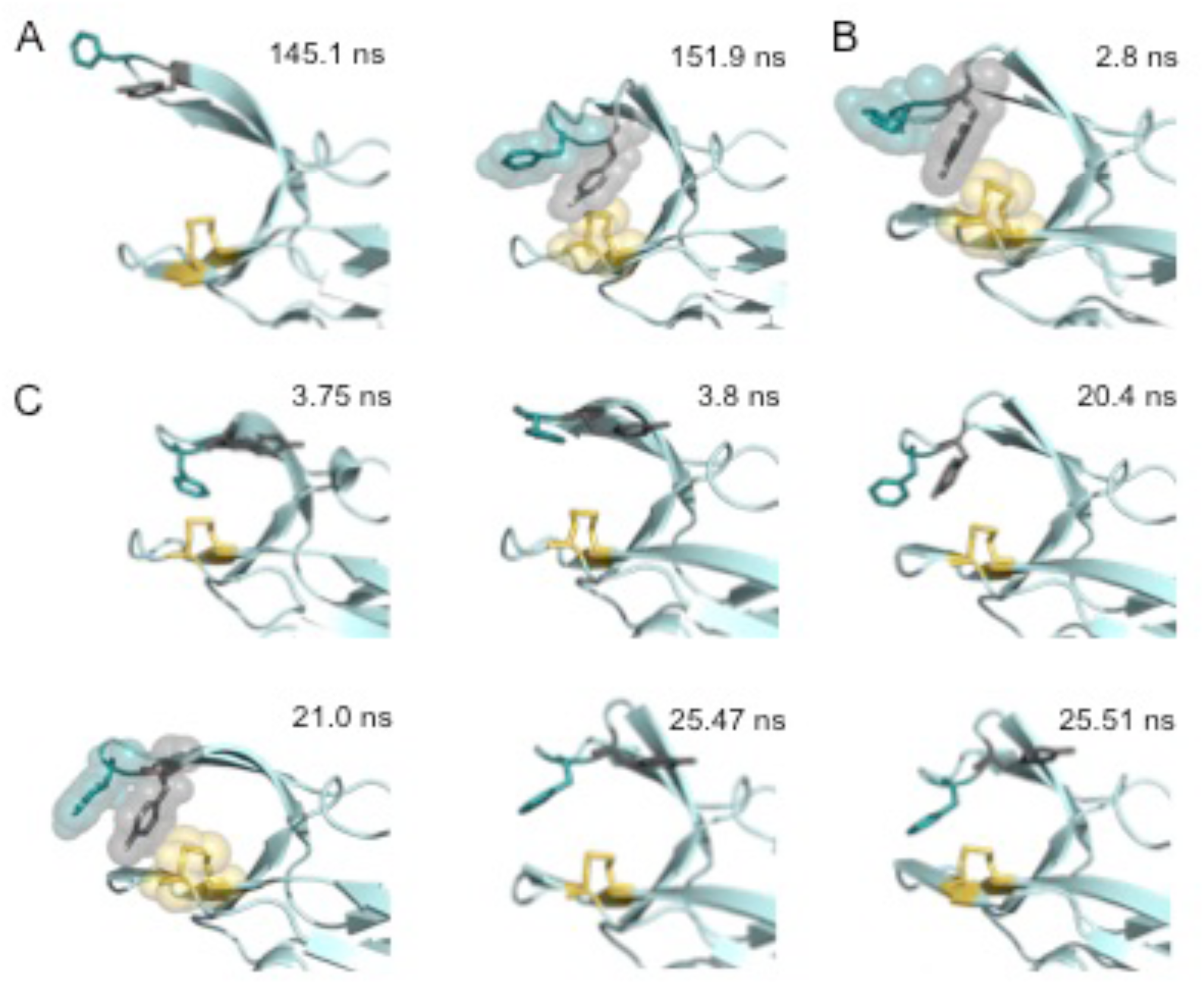
Snapshots of structures generated from the molecular dynamics simulations of nDsbD_ox_. (A) Two structures taken from the major loop opening described in Figures 5 and Figure 5 – figure supplements 1, 2 and 3. At 145.1 ns the cap loop adopts a very open conformation (F70-to-C109 distance of > 16 Å) that may be important for the initial docking of cDsbD leading to complex formation. At 151.9 ns Y71 adopts a position between F70 and C109 that blocks loop closing. (B) A structure taken from the loop closing simulation of nDsbD_ox_ shown in Figure 3G (in pink). At 2.8 ns Y71 adopts a position in between F70 and C109 which blocks the loop from closing immediately. (C) Six structures taken from the ∼22 ns loop opening event described in Figure 5 – figure supplement 4. Prior to loop opening, at 3.75 ns, F70 and Y71 both adopt the trans χ1 conformation with Y71 farther away from C109 than observed in the closed 1L6P X-ray structure. Loop opening at 3.8 ns involves F70 moving up and away from C109. At 20.4 ns the Y71 χ1 changes to the gauche-conformer bringing the ring of Y71 closer to C109 allowing it to adopt a ‘blocking’ conformation at 21 ns. Prior to loop closing Y71 returns to the trans conformation (25.47 ns) allowing F70 to approach C109 and leading to loop closure (25.51 ns). In all panels the backbone of nDsbD is shown as a cartoon in pale cyan. F70, Y71 and C103/C109 are shown in a stick representation and coloured deep teal, grey and yellow, respectively. A surface representation is used in the snapshots that show the ‘blocking’ conformation of Y71.

**Figure 5 – figure supplement 6.**
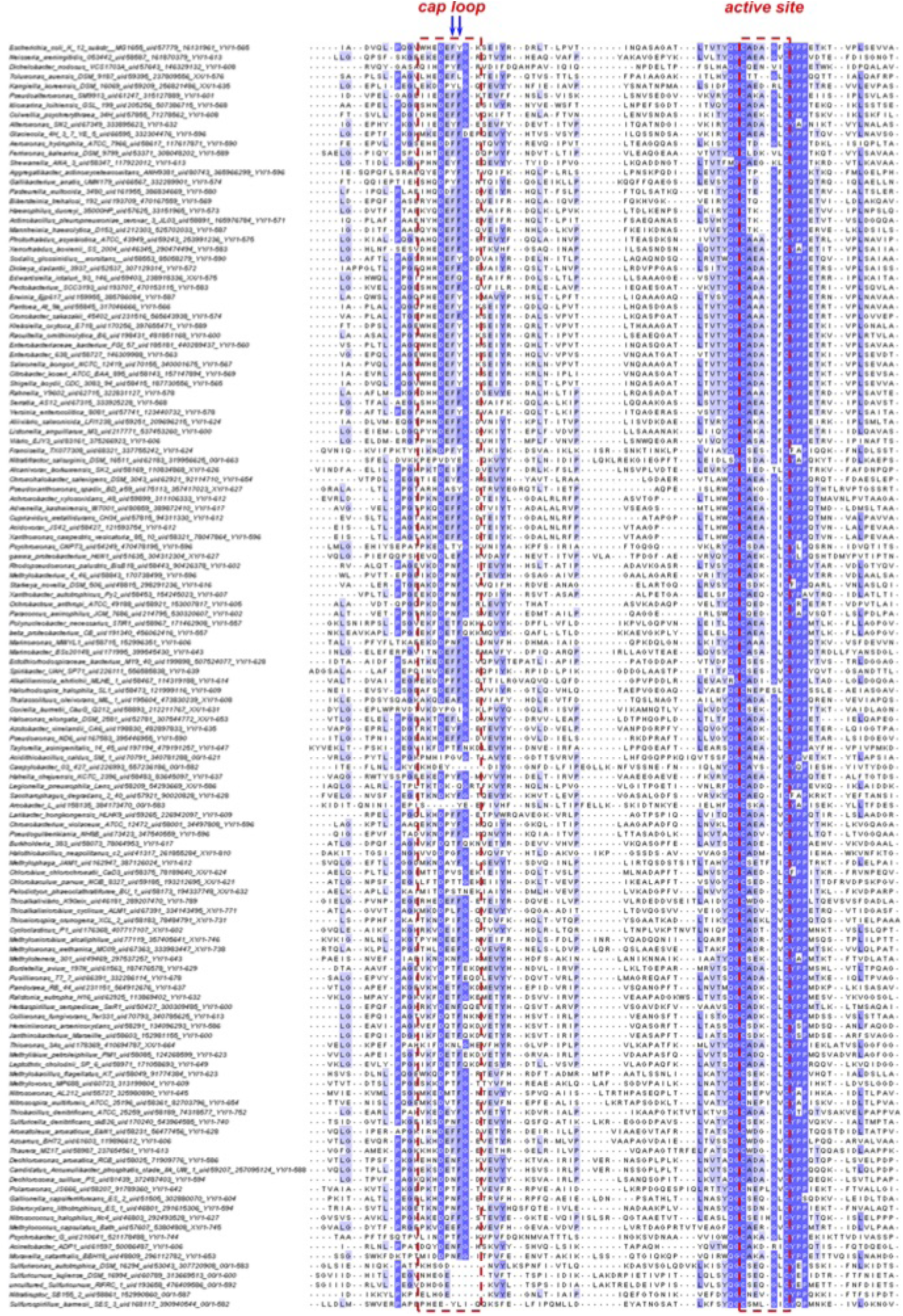
Sequence alignment of the cap-loop and active-site regions of nDsbD from β- and γ-proteobacteria. The figure shows a sequence alignment of representative nDsbD protein sequences from β- and γ-proteobacteria; only part of each nDsbD sequence is shown, spanning residues A53 to L119 in *E. coli* nDsbD. 2753 representative bacterial and archaeal proteomes were selected for homologues of nDsbD, and a subset of 731 sequences clustering around the *E. coli* DsbD were selected for subsequent analysis as a consistent phylogenetic group of proteins, belonging to mostly β- and γ-proteobacteria. 134 sequences, on per bacterial genus, were aligned along with *E. coli* nDsbD (top). The cap loop region and active-site cysteines are indicated by the red dashed lines, and positions 70 and 71 in *E. coli* nDsbD are indicated by blue arrows.

**Figure 6 – figure supplement 1.**
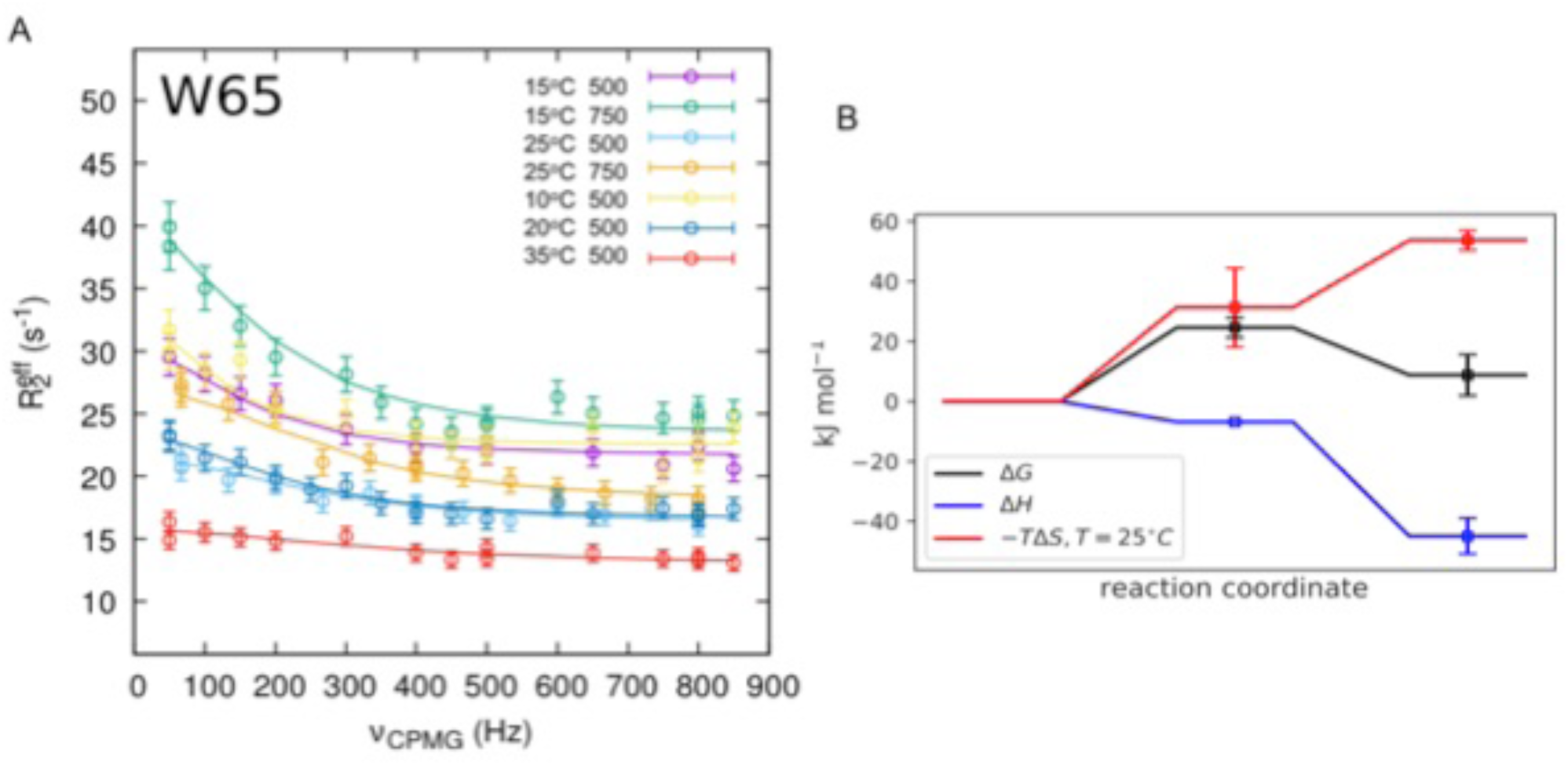
Analysis of the relaxation dispersion experiments at multiple temperatures. Global fitting of relaxation dispersion data measured for twelve residues in nDsbD_ox_ at 10 °C, 15 °C, 20 °C, 25 °C and 35 °C at 500 MHz and at 15 °C and 25 °C at 750 MHz was performed using CATIA (Cpmg, Anti-trosy, and Trosy Intelligent Analysis)(Alderson et al. 2019, Baldwin, Hilton, et al. 2011, Vallurupalli, Hansen, and Kay 2008), where the specific pulse sequence was simulated incorporating differential relaxation rates between the coherences generated, and off-resonance effects of the 180° pulses. Error bars were determined as described in Figure 6 – source data 1. Relaxation dispersion data measured for W65 and the curves from the global fitting are shown in (A) (Figure 6 – source data 1). The global fits at five different temperatures yield a description of the free energy landscape of the exchange process shown in (B). The minor state of nDsbD_ox_ is enthalpically more favourable than the ground state. This is reflected in the decrease in the population of the minor state from 7.1% to 1.6% when the temperature is increased from 10 °C to 35 °C (Figure 6 – source data 1). The minor state is entropically less favourable than the ground state. Overall, the reaction coordinate in (B) shows that the minor state is less favourable by 8.8 kJ mol^-1^ at 25 °C. The fitting of the experimental curves at the five temperatures also reports on the barrier between the two states. The transition state is enthalphically more favourable than the ground state. There is, however, a sizeable free energy barrier as the transition state is lower in entropy than the ground state, and so the rate for the forward reaction, while approximately temperature independent, decreases slightly with increasing temperature. Note that Δ*S*^‡^ and hence Δ*G*^‡^ depend on the transmission coefficient κ that is used in eqn. 1.

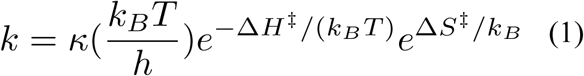 Δ*H*^‡^ and Δ*S*^‡^ are the activation enthalpy and entropy for the reaction. The transmission coefficient κ was set to 1.4 × 10^−7^ s^-1^ K^-1^.(Fersht 1984)

**Figure 6 – figure supplement 2.**
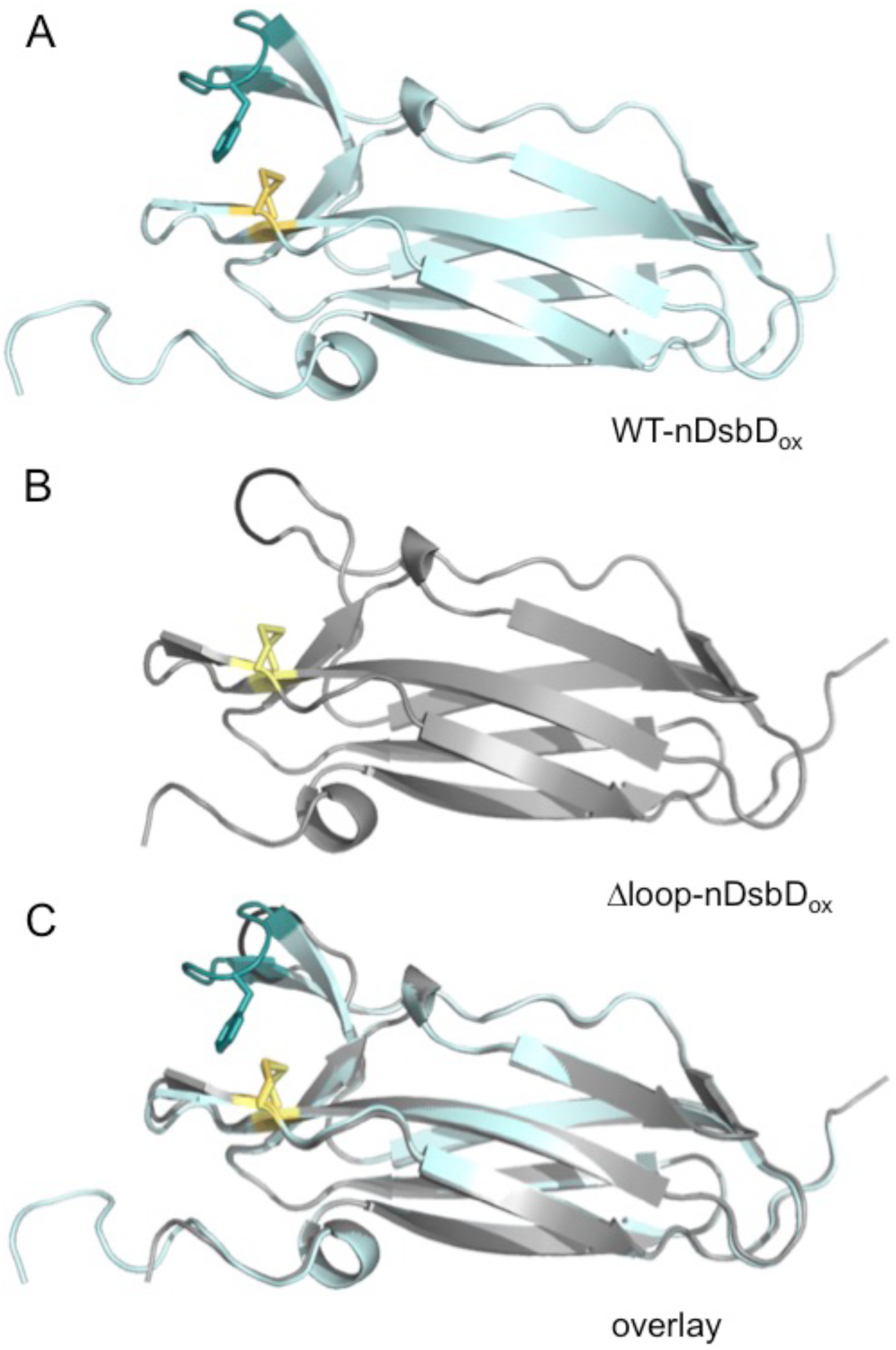
Comparison of the X-ray structures of wild-type nDsbD_ox_ (A) and Δloop-nDsbD_ox_ (B). The backbone/cap loop of wild-type nDsbD_ox_ (1L6P) and Δloop-nDsbD_ox_ (5NHI) are shown in cyan/deep teal and grey/black, respectively. F70 (dark teal), in wild-type DsbD_ox_, and the C103-C109 and C98-C104 disulfide bonds (yellow/pale yellow) are shown with a stick representation. The overlaid structures (C) have a Cα RMSD of 0.25 Å for residues 12-122.

